# PepSeek: Universal Functional Peptide Discovery with Cooperation Between Specialized Deep Learning Models and Large Language Model

**DOI:** 10.1101/2025.04.29.641945

**Authors:** Haifan Gong, Yue Wang, Qingzhou Kong, Xiaojuan Li, Lixiang Li, Boyao Wan, Yinuo Zhao, Jinghui Zhang, Guanqi Chen, Jiaxin Chen, Yanbo Yu, Xiaoyun Yang, Xiuli Zuo, Yanqing Li

**Affiliations:** Department of Gastroenterology, Qilu Hospital of Shandong University, Jinan, China; The Chinese University of Hong Kong, Shenzhen, China; Shandong Provincial Microecological Research and Biotherapy Center, Jinan, China; Shandong Provincial Clinical Research Center for Digestive Disease, Jinan, China; The Hong Kong University, Pokfulam, China

**Author notes:** These authors contributed equally to this work (Haifan Gong, Yue Wang, Qingzhou Kong, Xiaojuan Li).

## Abstract

Recent computational foundation models have revitalized the scientific discovery pipeline. However, developing foundational models for functional peptide discovery is costly due to the scarcity of wet-lab validated data. Meanwhile, conventional deep learning models are hard to generalize to unseen tasks or data distribution. Here, we introduce PepSeek, a universal approach for peptide discovery that synergistically integrates the most advanced large language model (LLM) with specialized small models. PepSeek harnesses the robust reasoning and generalization capabilities of LLM while leveraging the high predictive accuracy of specialized models trained for tasks such as antimicrobial activity regression and functional peptide generation. We have devised multiple collaborative strategies and task-specific modules demonstrating leading performance in peptide identification and generation. Notably, PepSeek achieves remarkable zero-shot prediction accuracy for peptides with diverse functionalities. We used PepSeek to identify a group of broad-spectrum antimicrobial peptide that exhibits low toxicity and high activity against drug-resistant bacteria, with the best surpassing all peptides currently undergoing clinical trials. Our framework establishes a new pipeline for scientific discovery with the help of LLM and specialized models.

## Introduction

In recent years, the field of computational biology has witnessed a surge of research centered around foundational models [1–7] and large language models (LLMs) [8– 14]. For foundational models, the typical research paradigm involves data collection, training large-parameter network models, and fine-tuning the models for downstream tasks [1, 2, 4, 15]. In the meantime, applying LLMs for research relies more on setting appropriate prompts [13, 16] or utilizing their internal embeddings [14] to represent the feature vector of single-cell data.

However, in the field of functional peptide design, the amount of high activity wet-lab validated peptide is still limited (less than 30,000) [17] compared with the billion-level data scale in genomic [4] (Fig. 1(e)) or medical imaging field [1, 15]. The limited high-quality wet-lab valid data has made it challenging to develop the foundation model for functional peptide discovery according to the scaling raw [18]. Besides, creating the foundation model is also costly and laborious [19]. For the identification tasks, the previous works [20–22] mainly focused on the development of lightweight machine learning models like random forests [23, 24] or neural networks [25, 26] for peptide discovery. These data-driven models take the limited wet-lab data as the supervision, which can easily face performance degradation in the unseen data distribution or task [27, 28]. The performance degradation is called the “out-of-domain” issue in machine learning [29], where there are two typical situations in functional peptide discovery: one is the peptide sequence similarity gap between the training set and testing set (e.g., testing set with the peptide sequence similarity to training set *<*30%), another is the task gap, which indicates the training set focus on antimicrobial, but the target tasks focus other tasks like anti-virus or anti-fungus. We are thus prompted to examine whether LLMs [30, 31] might be helpful to enhance the generalization performance of conventional deep learning models by exploiting their intrinsic capacity for sample-independent reasoning thus prompted to examine whether LLMs [30, 31] might be helpful to enhance the generalization performance of conventional deep learning models by exploiting their intrinsic capacity for sample-independent reasoning ability [32].

**Fig. 1.**
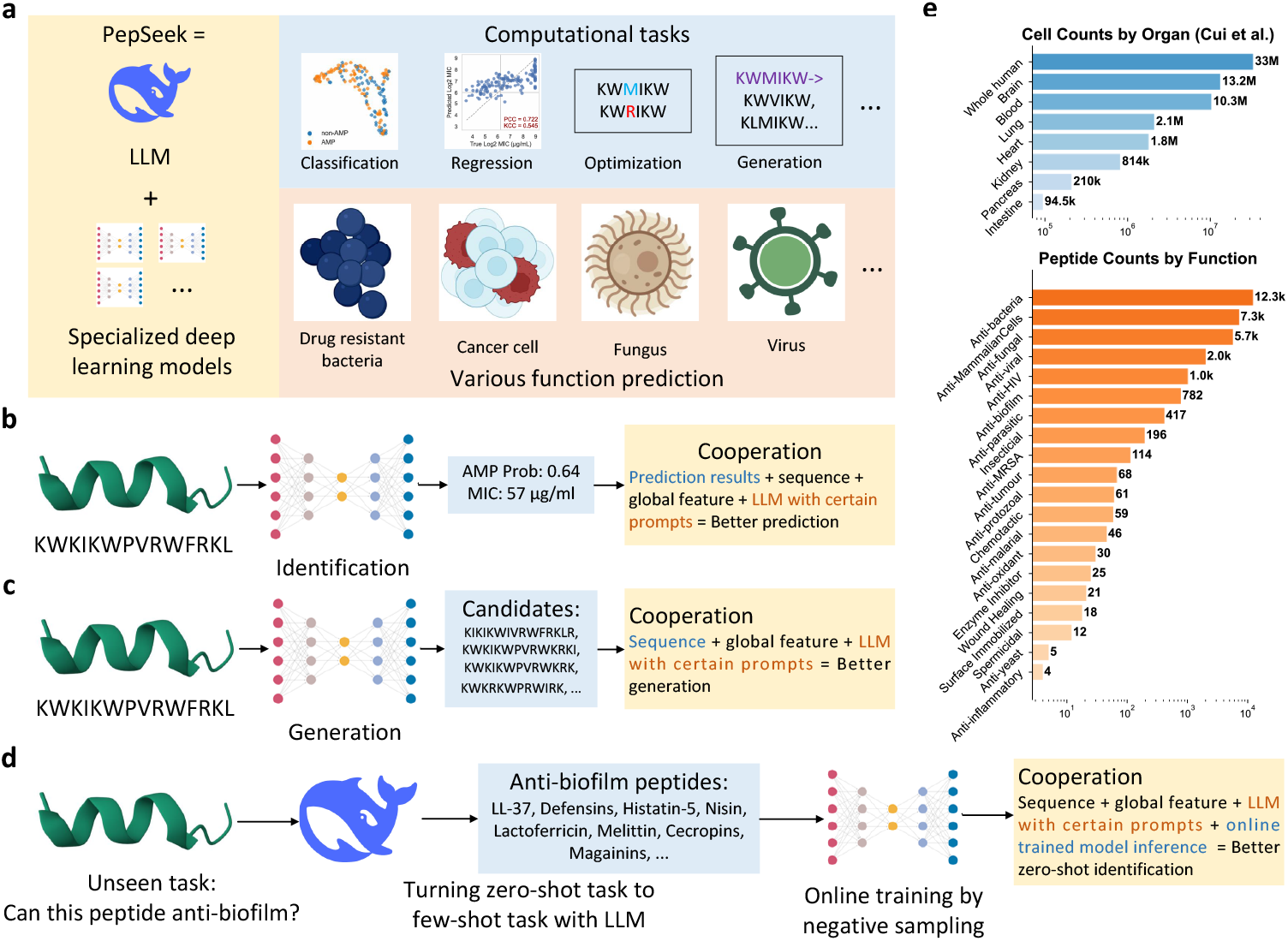
An overview of the semantic paradigm of our work. **a**. PepSeek leverages the generalization ability of large language models (LLMs) in conjunction with specialized deep learning models to address various computational tasks and functional prediction challenges. **b**. For peptide identification, specialized predictions from deep learning models are further refined by the LLM. **c**. In peptide generation, a reinforcement learning model first proposes sequences that the LLM then improves to enhance activity and lower toxicity. **d**. In the zero-shot peptide identification scenario, the LLM gathers peptides with specific functions, which are subsequently used in a deep-learning model for detailed functional prediction. **e**. The figure shows that peptide data are far more limited compared to single-cell data, thereby limiting the potential for training a foundation model.

With the rapid development of natural language processing technologies [8], LLMs [9, 30, 33] excel in expressive capabilities and generalization performance, especially with their reasoning ability [31, 34] by chain-of-thoughts [32] training pipeline. Since these LLMs are trained based on human knowledge [9], it is feasible to partly address the out-of-domain issue using their outstanding generalization and contextual understanding ability. Still, LLMs have several drawbacks, such as being inadequate for data-driven tasks like predicting specific minimum inhibitory concentration (MIC) values. Besides, LLM’s hallucination [35, 36] can lead to unreliable results. The above phenomenons make us wonder if we can use the input peptide’s global descriptions and the conventional small deep learning model’s output to reduce the likelihood of model hallucinations, enhancing the reliability and accuracy of the results.

Based on the above observation and consideration, we propose PepSeek (Fig. 1a), a cooperation framework between LLMs and specialized expert models to achieve efficient, reliable, and flexible peptide design. The intuition of PepSeek is leveraging the strengths of LLMs while compensating for the shortcomings of the specialized small models. Specifically, we first collect existing peptide data and train a series of specialized deep-learning-based models for peptide identification and generation. With the trained specialized small models, we further design several simple yet effective cooperation pipelines between specialized deep learning models and LLMs: In the peptide identification process in Fig. 1b, predictions derived from specialized models undergo refinement by the LLM, ensuring higher accuracy. For peptide generation in Fig. 1c, a reinforcement learning model initially generates candidate sequences that the LLM further optimizes to achieve enhanced biological activity and reduced toxicity. In the zero-shot identification setting (Fig. 1d), the LLM is employed to curate peptides of specific functional and then used to train a specialized deep learning model; after that, we use the cooperation pipeline for further function identification. By taking advantage of the reasoning ability of LLMs, we can better identify or generate the peptides and know how the peptide is determined or selected with the transparent reasoning process [31] (Fig. 6). We evaluated our PepSeek on the peptide identification (classification and MIC regression tasks) and generation (unconditional generation and template-based optimization) tasks. The results have shown that PepSeek can significantly boost the identification performance of peptides on the out-of-domain data and generate more reliable peptides. Moreover, the out-of-domain function prediction task shows PepSeek can make zero-shot predictions accurately on 11 unseen functions, as shown in Fig. 5. Our findings show that the cooperation is more effective than previous approaches.

To sum up, we have made the following contributions to this work: (1) A new application paradigm of AI models in computational biology is proposed by implementing a straightforward collaborative approach between specialized deep learning models and LLMs. PepSeek provides a new solution for peptide design and offers excellent scalability, allowing easy extension to other areas within computational biology. (2) PepSeek shows unique advancements in the out-of-domain task for out-distribution identification and functional peptide discovery. (3) Through extensive computational and wet-lab experiments, PepSeek has demonstrated exceptional performance. The peptides identified exhibit significantly better efficacy in inhibiting bacteria than all existing peptides with higher activity and lower toxicity. The success of PepSeek not only advances peptide design technology but also introduces more efficient and intelligent tools to the field of computational biology, offering broad application prospects and profound impacts.

## Results

### Cooperation between LLM and Specialized Deep Learning Model for Peptide Classification and Regression

Despite the discovery of over ten thousand functional peptides with potential applications in combating species like drug-resistant bacteria [17, 37], the number of peptides with both high activity and low toxicity remains limited [17]. Unlike previous studies that solely dichotomized peptides into bioactive and non-bioactive categories, we followed the methodology described in [38] by categorizing peptides based on their minimum inhibitory concentration (MIC). Specifically, peptides with an average MIC to bacteria below 128 were considered active. Considering the scarcity of peptides with less common functions, such as antiviral activity, we focused on the more prevalent anti-bacterial activity for training and evaluating our specialized model.

For the training and validation sets, data were compiled from DBASSP [39], QLAPD [38], with additional negative samples from UniProt (http://www.uniprot.org) by assigning them a MIC value of 512. Five-fold cross-validation was conducted on this dataset, and the model was additionally tested on the QLX test set with 151 synthetic peptide sequences [38] that are valid in our web lab. The private QLX test set provides a unique advantage in comparing the LLM-based methods since the LLM-based methods use the data from the internet and manuscripts for pre-training. There is no potential data leakage issue with this private set. To avoid potential data leakage and better simulate the out-of-domain situation, peptide sequences with a maximum internal similarity exceeding 30% (as measured by the Needleman–Wunsch algorithm [40]) were removed between the training/validation set and the test set.

All models were trained for 120 epochs, and test performance was determined by averaging the performance metrics from the final epoch across the five folds. For the classification task, evaluation metrics included the area under the precision-recall curves (AUPRC) and the area under the receiver operating characteristic curve (AUROC) [22]. For regression, mean absolute error (MAE), relative squared error (RSE), Pearson correlation coefficient (PCC), and Kendall correlation coefficient (KCC) were employed for a more comprehensive comparison [41]. We use the torchmetrics [42] for the implementation of the metric.

To prevent potential overfitting associated with models possessing a large number of parameters, such as ProGen [19, 43], we chose to use lightweight models following the previous work [21, 22], including Mamba [44] and Multi-head Latent Attention (MLA) [45], as shown in Fig. 2a. These were evaluated alongside conventional architectures such as LSTM [25], Transformer [26], and Multi-head Attention (MHA) [26] for peptide classification and regression. Unlike earlier works [20, 22, 46] that employed a straightforward ensemble of various models, we introduce the LLM-guided model ensemble (LLM-GME) (Fig. 2b), which use the LLM to select models based on their validation performance with the prompt in Fig. 2c. Our results indicate that the current LLM, with its inherent reasoning capabilities, can identify a minimal subset of models that achieves comparable or superior performance.

**Fig. 2.**
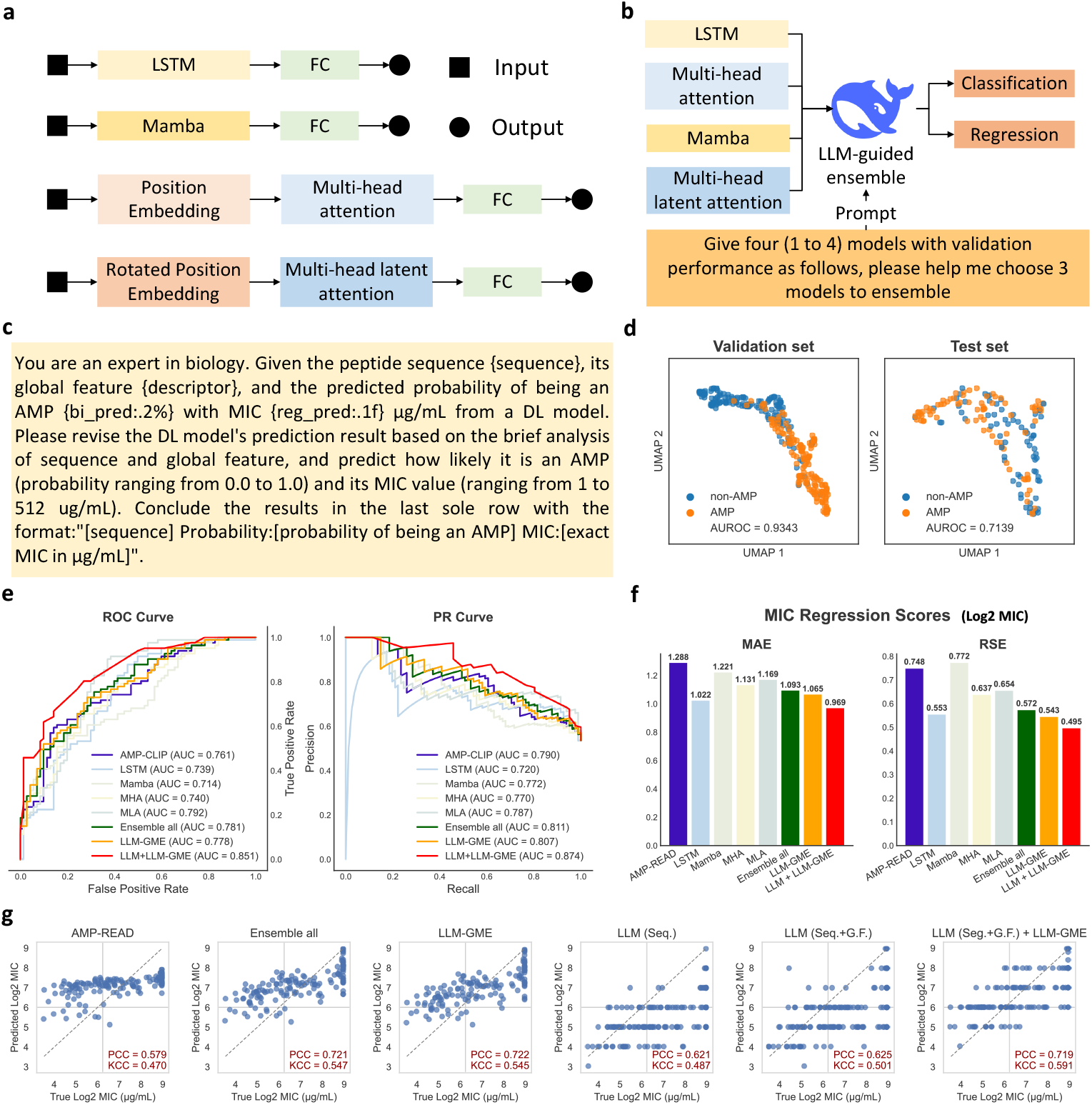
Overview of the design and performance of PepSeek for peptide identification tasks. **a**. The detailed architecture of the specialized deep learning models. **b**. The LLM-guided model ensemble (LLM-GME) for classification and regression tasks. **c**. The prompt facilitates the cooperation between the LLM and LLM-GME. **d**. A 2D UMAP visualization of the Mamba model’s feature representations for peptide classification on the in-domain validation set and out-of-domain testing set, indicating significant performance degradation with a more confusing decision boundary. **e**. ROC and PR curves comparing current state-of-the-art methods with our approach to the classification task. The results suggest that modern network structures like MLA and Mamba enhance ensemble performance. Although the LLM-guided ensemble slightly lowers classification performance, it substantially reduces the overall parameter count. **f**. MIC regression results evaluated by MAE and RSE metrics, demonstrating that the LLM-guided ensemble achieves improved regression performance with fewer parameters compared to ensembles that directly combine all models. **g**. PCC and KCC scores for the MIC regression task, showing that introducing global features enhances the LLM’s regression capability by yielding a comparable PCC score and a significantly improved KCC score relative to using only small models.

Still, when applied to the unseen data, the out-of-domain issue will hurt the practicality of the above small model [47]. To valid this, we present a 2D UMAP visualization in Fig. 2d, which compares the feature representations from the Mamba model across in-domain (validation) and out-of-domain (testing) datasets. A more confused decision boundary is observed in the out-of-domain dataset with the decreased AUROC from 0.93 to 0.71. To alleviate the performance degradation due to the out-of-domain issue, a cooperation pipeline is established by using the output of the specialized model as a prompt to the LLM [31], along with the original peptide sequence and global physicochemical property features. This integration, capitalizing on the LLM’s reasoning ability, enables more precise and interpretable predictions in anti-bacterial classification and regression tasks. From the results shown in Fig. 2e,f,g, we can observe that the cooperation between LLM and LLM-selected specialized small deep learning models can significantly boost the AUROC and AUPRC while reducing the regression error.

Furthermore, Fig. 2e presents ROC and PR curves that benchmark our approach against state-of-the-art methods, indicating that structures such as MLA and Mamba, when employed in tandem with LLM guidance, achieve a competitive balance between performance and model complexity. Finally, the panels Fig. 2f and Fig. 2g highlight the advantages of our proposed method in the context of MIC regression tasks; the LLM-guided ensemble not only reduces the model parameter count but also improves regression metrics, evidenced by lower MAE and RSE values and enhanced PCC and KCC scores when compared to traditional small model setups.

We benchmarked several state-of-the-art LLMs, including GPT-4o, o3-mini, DeepSeek-V3 and DeepSeek-R1 in Fig. 3. This experiment unveils the identification of discrepancies between reasoning and non-reasoning types and between vendors OpenAI and DeepSeek, while also test the stability of model’s prediction.

**Fig. 3.**
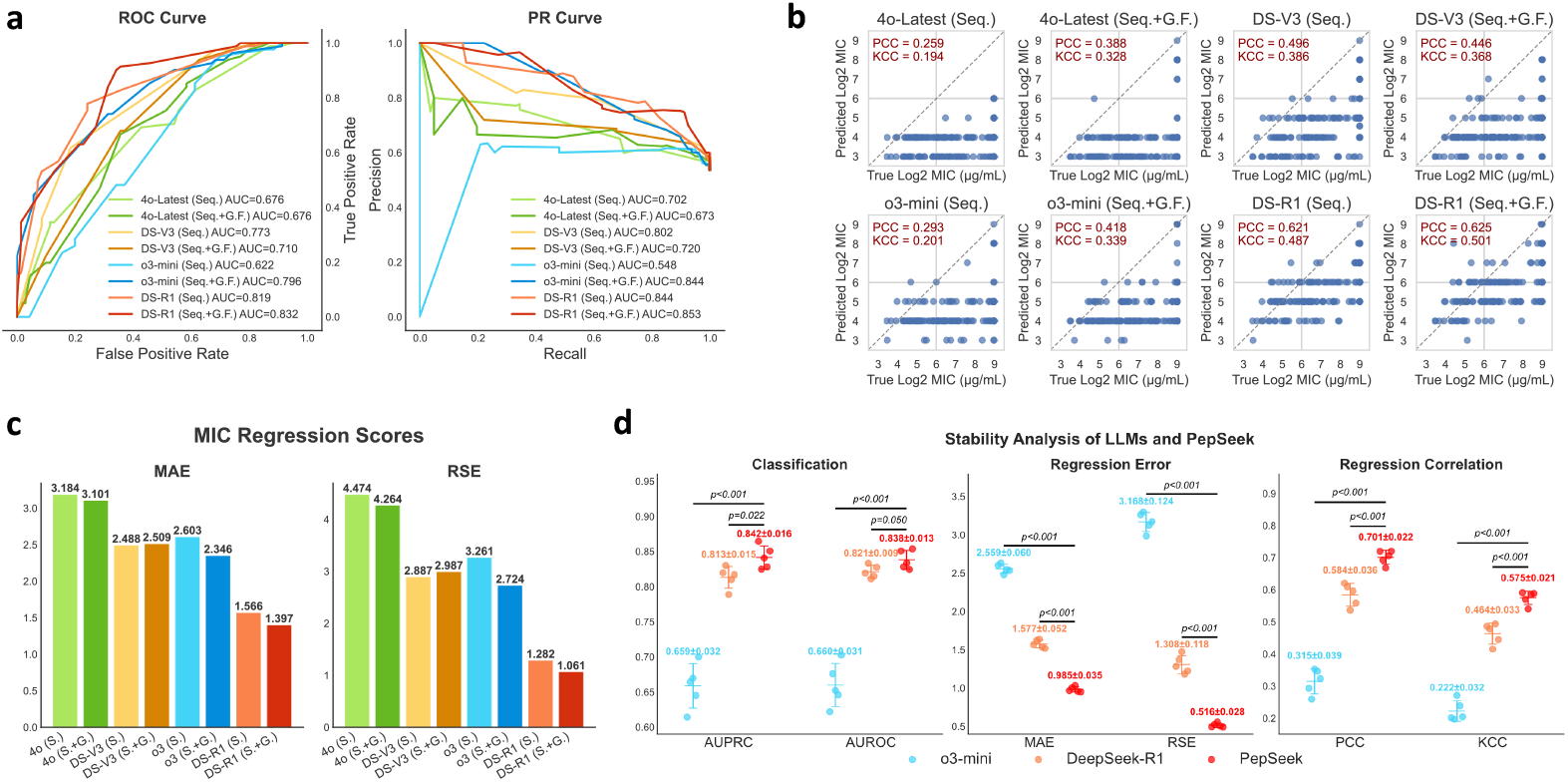
Comparison of multiple LLMs and the proposed method. **a**. ROC and PR curves for state-of-the-art LLMs. The results indicate that reasoning LLMs achieve superior classification performance relative to non-reasoning models. Moreover, incorporating global features enhances the performance of reasoning LLMs while slightly impairing that of non-reasoning variants. **b**. PCC and KCC scores for the MIC regression task. The findings demonstrate that DeepSeek models outperform OpenAI models and that adding global features leads to an overall improvement in performance. **c**. MAE and RSE scores for the MIC regression task. DeepSeek models continue to surpass OpenAI models, with DeepSeek-R1 exhibiting a significant performance lead. **d**. Stability analysis of OpenAI o3-mini, DeepSeek-R1, and the proposed method PepSeek. The analysis reveals that o3-mini is the least stable and performs the poorest, whereas PepSeek is the most stable and accomplishes a marked advantage in MIC regression performance.

Fig. 3a presents the ROC and PR curves for classification performance. Incorporating global features improves the performance of reasoning models but degrades that of non-reasoning models. For instance, GPT-4o’s AUPRC drops from 0.702 to 0.673, while o3-mini’s increases from 0.548 to 0.844. DeepSeek models show similar trends, with V3 falling from 0.802 to 0.720 and R1 slightly rising from 0.844 to 0.853. We attribute this to non-reasoning LLMs tending to overlook sequence-specific details in favor of global features, unlike reasoning LLMs that leverage all available prompt information in their chain-of-thought reasoning process. Additionally, DeepSeek models significantly outperform OpenAI models, likely because the training data for the latter contains less information on antimicrobial peptides.

Fig. 3b and c show the MIC regression performance. Reasoning models generally outperform non-reasoning ones, and DeepSeek models significantly exceed OpenAI models, which is consistent with the classification results. Incorporating global features can enhance performance, although DeepSeek-V3 experiences a slight decline. Notably, DeepSeek-R1 demonstrates a substantial lead over the other models.

As demonstrated in Fig. 3d, the stability of the proposed method PepSeek was evaluated by conducting a comparative experiment with OpenAI’s o3-mini, DeepSeek-R1, and the aforementioned method. Each of these approaches was subjected to five distinct trials. The results indicated that o3-mini exhibited substandard performance and stability compared to the other two methods. Conversely, PepSeek demonstrated superior performance and stability, with all six classification and regression metrics having *p*-values no more significant than 0.005 and a particularly noteworthy and consistent regression performance.

### Cooperation between LLM and Prior-Guided Reinforcement Learning for Peptide Generation and Optimization

Another challenge for peptide design is to rapidly and cost-effectively generate or optimize peptide sequences *in silico* rapidly and cost-effectively. To address this, we developed an online prior-guided reinforcement learning (PRL) agent for peptide optimization by taking advantage of domain knowledge, where the pipeline is in Fig. 4a. We reformulated peptide generation as a reinforcement learning [48] task (Fig. 4b): the template peptide sequence defines the state space, amino acid mutations represent the action space, and the desired physicochemical properties constitute the reward signal.

**Fig. 4.**
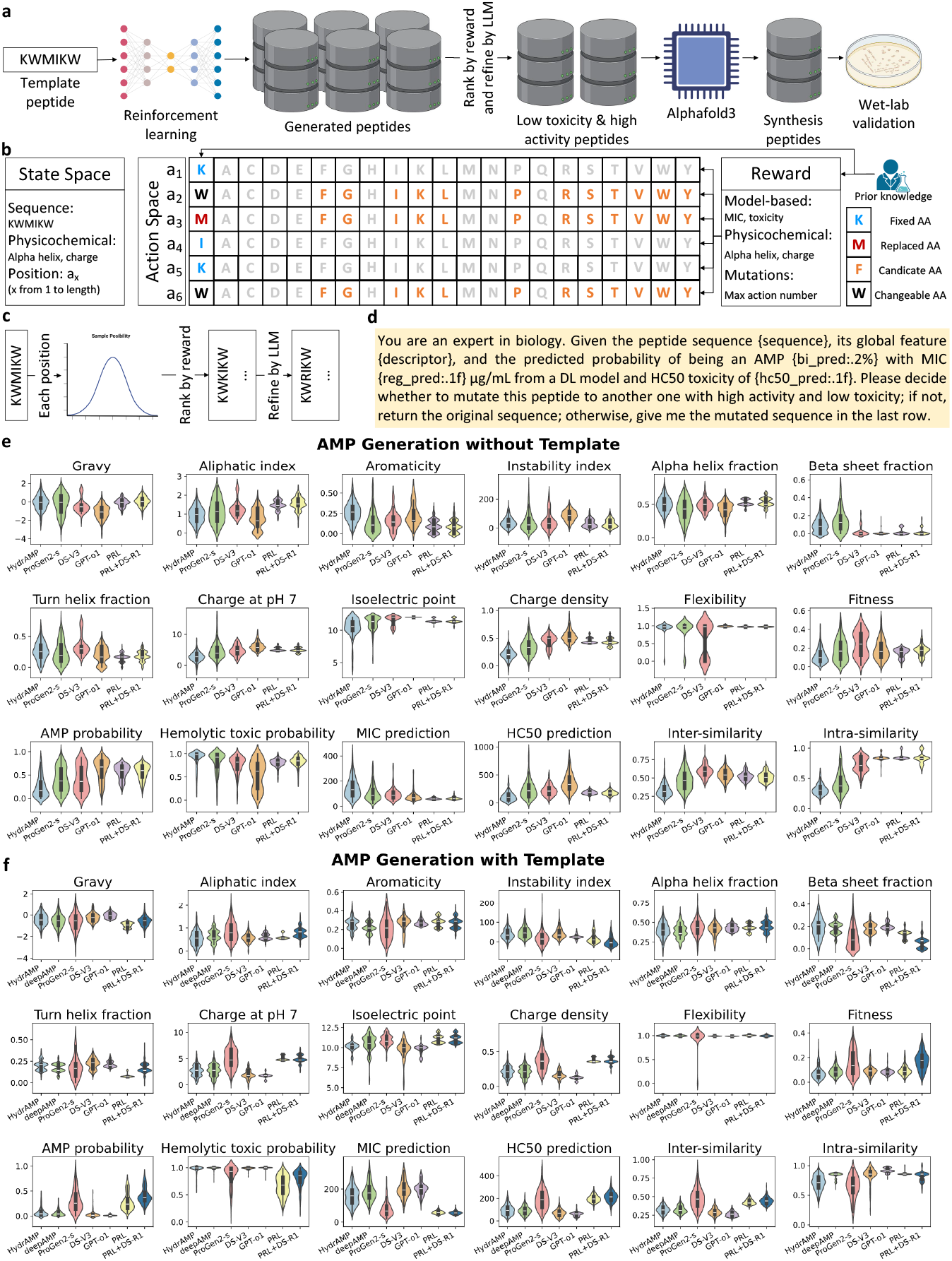
Model architecture and performance for peptide generation. **a**. The pipeline for peptide optimization to generate candidate peptides. **b**. design of the prior-guided reinforcement learning (PRL) leverages prior knowledge of potential candidate amino acids to narrow the search space and incorporates physicochemical features for further refinement. **c**. The inference phase for peptide optimization using PRL. **d**. The prompts used during peptide optimization with the LLM. **e**. Comparison between our method and state-of-the-art approaches for AMP generation (sample size of 200). The results demonstrate that peptides solely generated by the LLM are unstable and exhibit low toxicity. In contrast, the cooperation between PRL and DS-R1 enhances both the effectiveness and diversity of the generated peptides. **f**. Analysis for template-based AMP generation (sample size of 200), showing that our PRL effectively optimizes the given peptide sequences to achieve improved physicochemical properties such as better fitness and reduced instability, while the cooperation slightly increases activity at the expense of a marginal rise in toxicity.

**Fig. 5.**
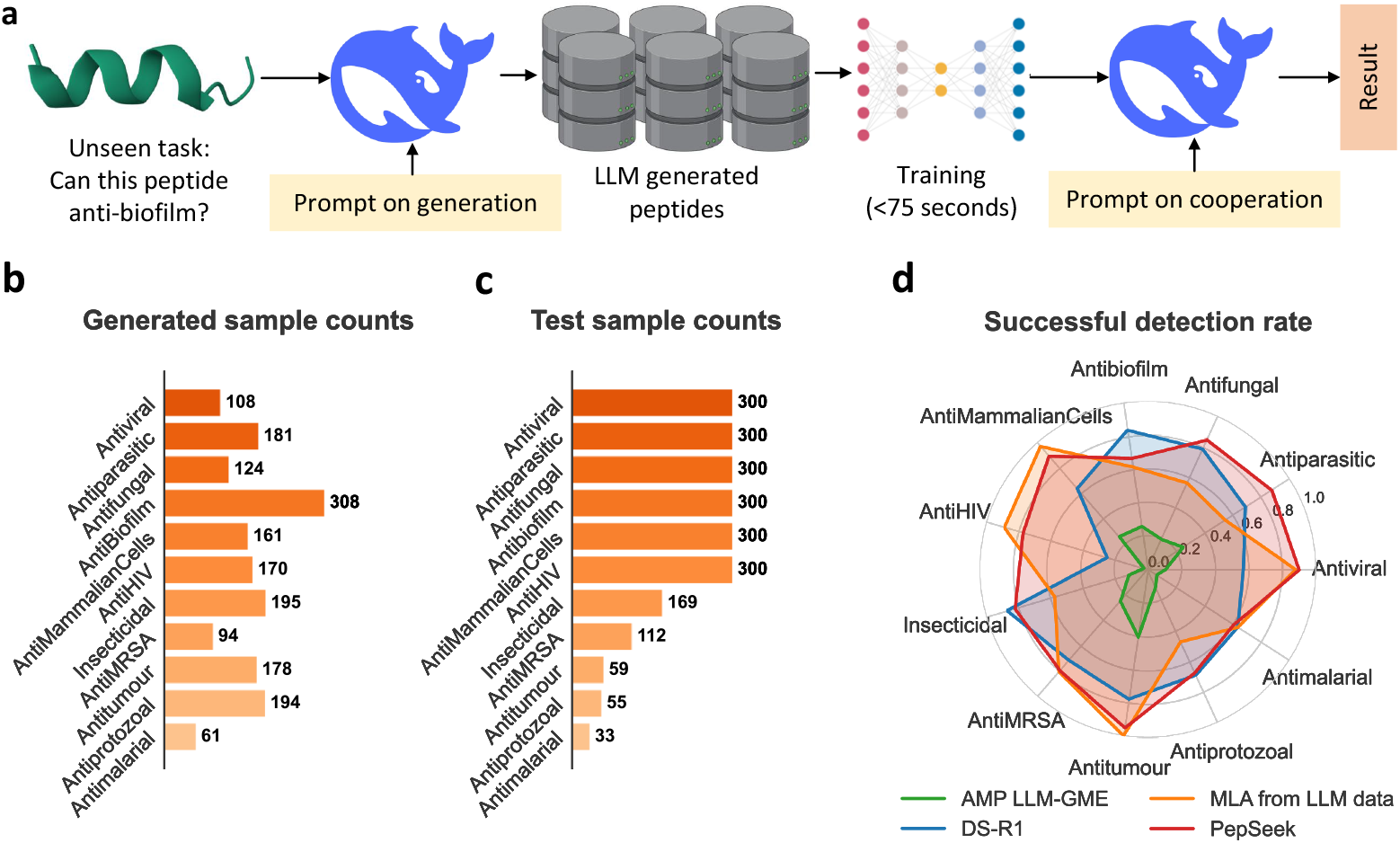
Overview of Zero-Shot Prediction Tasks. **a**. The pipeline that transforms a zero-shot prediction task into a few-shot prediction task. **b**. The data distribution of LLM generated positive samples after eliminating duplicates and ones already in the test set. **c**. The data distribution of the test set for our zero-shot prediction task. **d**. Performance comparison between the LLM’s direct predictions and the cooperative prediction results, measured by the successful detection rate. The results indicate that the additional reasoning ability provided by the cooperative approach can better clarify peptides with certain functions and that generating peptide sequences via the LLM followed by training a specialized deep learning model leads to significantly improved performance.

**Fig. 6.**
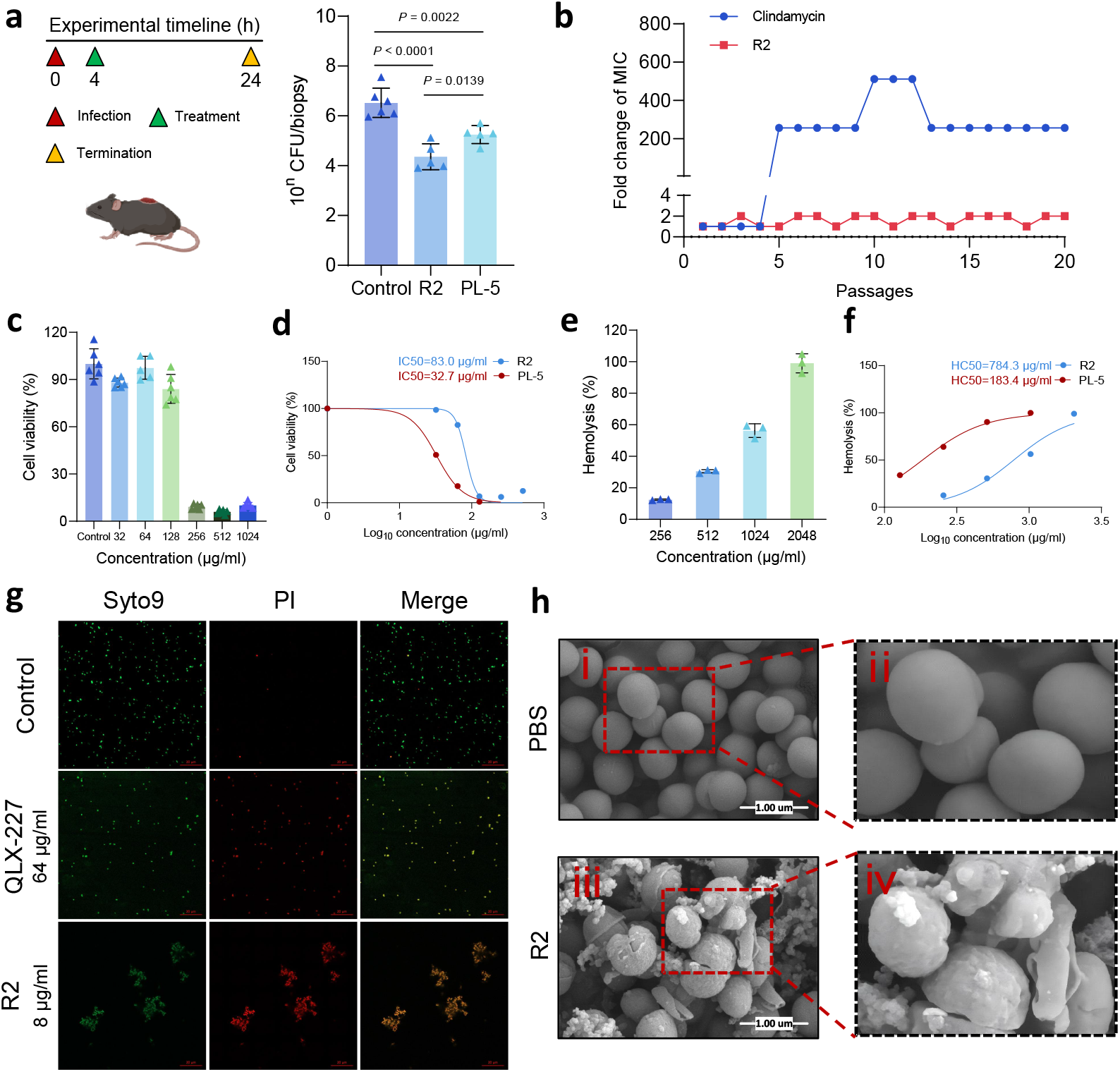
Overview of the Biological Properties of R2 and Its Antibacterial Activity. **a**. In vivo antibacterial efficacy of R2 in a skin infection model, demonstrating superior performance compared to the potent antimicrobial peptide PL-5 (Peceleganan). **b**. Assessment of resistance acquisition in *Staphylococcus aureus* (S. aureus CMCC 26003) cultured with clindamycin at sub-minimum inhibitory concentration (1/2× MIC, panel g; n = 3, biologically independent replicates). In vitro evaluation of R2’s biological properties, including cytotoxicity on mouse embryonic fibroblast cells (NIH3T3, panel c) and hemolytic activity (panel e), with mean ± standard deviation data. Determination of the 50% inhibitory concentration (IC_50_ on NIH3T3 cells, panel d) and the 50% hemolytic concentration (HC_50_, panel f) for both R2 and PL-5. **g**. Confocal laser scanning microscopy (CLSM) images (panel g) of S. aureus stained with the LIVE/DEAD BacLight kit after treatment with PBS, QLX-227-1, and R2 (scale bar, 20 *µ*m). Scanning electron microscopy (SEM) images (panel h) showing morphological changes in bacteria treated with PBS (panels h i-ii) and R2 at 1× MIC (panels h iii-iv), with a scale bar of 1 um.

Prior knowledge is embedded into both the action space and the reward function, as shown in Fig. 4b. For example, amino acids with negative charges, less favorable for enhancing performance, are masked in the action space. The reward function comprises two components. The step-wise reward captures metrics including the physicochemical properties like aliphatic index and overall charge. In contrast, after multiple mutation steps, the global reward evaluates the final optimized peptide, accounting for the number of mutations and the aggregate physicochemical properties. In addition to these static properties, rewards derived from deep learning models—specifically, the previously trained AMP identification, MIC regression, and toxicity classification models—are incorporated to guide the search toward superior sequences. The proposed PRL framework applies to peptides generated by conditional or unconditional generation models. The inference phase, depicted in Fig. 4c, demonstrates the practical application of PRL in generating optimized peptides.

We further established a cooperative paradigm between the specialized PRL and LLMs. The prompts shown in Fig. 4d exemplify the facilitation of this process. This scheme provides the generated peptide sequence as input to the PRL model. At the same time, the LLM is prompted to modify the peptide composition to enhance activity further and reduce toxicity. The combined approach yields peptide ensembles with improved functional attributes.

We evaluated the optimized peptides based on their physicochemical properties and *in silico* predictions with the best specialized deep learning models. Our results demonstrate that the PRL framework significantly outperforms previous methods in conditional and unconditional peptide generation tasks. The performance evaluation in Fig. 4e compares our approach against leading AMP generation methods, revealing that while LLM-only generated peptides tend to be unstable with low toxicity, the combined strategy of PRL and DS-R1 not only increases peptide effectiveness and diversity but also maintains a balance in toxicity. Furthermore, Fig. 4f presents a comparative study on template-based AMP generation, where the template is the sequence “KWMIKWPSNWFTML” from [38]. Our PRL successfully optimizes given peptide sequences to yield better physicochemical properties, such as enhanced fitness and decreased instability, alongside a noticeable improvement in activity albeit with a slight increase in toxicity. Overall, these results substantiate the efficacy of our approach in generating superior peptide candidates.

### Cooperation between LLM and Online-Trained Specialized Model for Zero-Shot Functional Peptide Discovery

The discovery of novel functional peptides is hindered by data scarcity. To overcome this limitation, we harnessed the capabilities of a powerful LLM to perform zeroshot peptide function prediction. Testing data were drawn from dbAMP 3.0 [37] and QLAPD [38], which comprises peptides with 19 distinct functions. A set of 11 functions was selected for a comprehensive evaluation by removing the functional peptides with a count of fewer than 30. This large testing sample size can provide a more meaningful analysis. Recognizing that peptides not present in this database may still possess specific functions, we evaluated our cooperative pipeline by treating all available samples as positive instances. We then assessed performance using the successful detection rate (SDR, %), defined as the fraction of peptides with a recognized function relative to the total number of functional peptides in each category.

As shown in Fig. 5a, PepSeek converts a zero-shot prediction task into a few-shot scenario with the LLM, enabling the small model to leverage a limited number of annotated examples for improved performance. Specifically, we initially prompted an LLM to generate peptides linked to the target function to address zero-shot functional discovery. After eliminating duplicates and peptides already present in the dbAMP database, as shown in Fig. 5**b**, the resulting peptides ranged from 61 to 308 samples per function and were designated as positive examples; meanwhile, 9,108 known non-AMP peptides from databases were used as negatives. These samples constituted the training dataset for the subsequent training of specialized, small-scale peptide classification models. Resampling techniques were employed to maintain a balanced positive-to-negative ratio during training. For validation, the sets comprised positive samples from the corresponding function in the dbAMP database and 49 non-AMPs as negative samples from the QLX test. The validation F1 score was adopted as the metric for selecting the optimal model weights, thus mitigating the risk that the model predicts all samples as positive.

Fig. 5c presents the data distribution of the test set utilized in these tasks, accentuating the challenges of varied and sparse data. Functions with more than 300 samples in the dbAMP database were reduced to 300 by random selection, thus preventing the LLM from taking too long to complete the task, resulting in 2228 samples for the test set.

Fig. 5d compares the performance in the successful detection rate of direct predictions made by the LLM with those achieved via our cooperative approach. In the cooperative pipeline, the peptide sequences generated by the LLM are first used to enhance the training of the MLA model. Subsequently, the MLA model makes its predictions, which are further refined by the LLM, leading to a significantly higher overall detection rate. This improvement highlights the importance of incorporating additional reasoning capabilities to identify peptides with the desired functions more effectively. Our experimental results show that a specialized deep learning model, trained solely on anti-bacterial peptides, struggles to recognize peptides with other functions (average SDR of 18.53%). In contrast, the LLM did better at directly identifying peptides with specific functional attributes from their sequences (average SDR of 67.83%). This is because the LLM’s training data included extensive details about peptides with diverse functions, effectively converting a zero-shot task into a few-shot prediction problem. By incorporating these generated samples into the specialized model’s training, the original out-of-domain challenge is successfully recast as an in-domain problem (average SDR of 72.3%). Finally, the PepSeek method further leverages the LLM’s knowledge by prompting it with both the specialized model’s results, the peptide sequence, and its global features, leading to an even better overall performance (average SDR of 80.11%).

In order to provide further validation of the usability of the method under investigation, a case study was conducted using two samples from the wet lab experiment. R2 and PL-5. As demonstrated in Tab. 2, both R2 and PL-5 exhibited mean MIC values for fungi and MRSA species of less than 128 *µg/mg*, and HC 50 for human cell is also less than 128 *µg/mg*, indicating their potential for antifungal, anti-MRSA, and antimammalian cell activities. As demonstrated in Tab. 2, PepSeek predicts R2 and PL-5 as active antifungal, anti-mammalian cell, and anti-MRSA agents, consistent with the wet lab results.

### In vitro and in vivo evaluation of Top 10 antimicrobial peptides candidate for efficacy

We use the pipeline detailed in Fig. 4a to optimize the peptide QLX-227-1 from our previous work [38]. We first train the PRL with the rewards functions (i.e., toxicity classification, activity classification, MIC regression). Then, we conduct the inference phase of the PRL model to generate the candidate peptides ranked by the reward. We further used the LLM to optimize the candidates. We chose the top 80 peptides. After that, we use Alphafold [49] to predict the 3d structure of the peptide. Then, we calculate the physicochemical properties and rank them by the alpha helix and stability to get the top 10 peptides for synthesis. We comprehensively evaluate the antimicrobial efficacy and potential economic viability for industrial drug production with the 10 antimicrobial peptides (R1–R10).

The antimicrobial activity of these peptides was assessed using standard bacterial strains and multidrug-resistant bacterial strains isolated from clinical settings covering the ESKAPE pathogens. Of all candidate peptides, R2 exhibited the lowest geometric mean minimum inhibitory concentration (MIC) (Table 1). R2 showed stronger antibacterial activity than the source peptide QLX-227-1. Further comparative analysis with several antimicrobial peptides (Peceleganan, C16G2, D2A21, Omiganan, OP-145, and DPK-060) [50] currently in clinical trials still revealed that R2 exhibited higher broad-spectrum antimicrobial activity against all tested superbugs (Table 2), demonstrating comparable or superior efficacy to the currently best-performing broad-spectrum AMP in clinical trial, PL-5 (Peceleganan) [51]. Following confirmation of *in vitro* antimicrobial activity, we established a full-thickness skin excision infection model in mice to evaluate its *in vivo* efficacy, with PL-5 as a comparator. Four hours after inoculating the wound sites with *S. aureus*, drug interventions were applied with the dosage of 0.1 mg. Tissue bacterial load analysis at 24 hours post-treatment showed that R2 sig-nificantly reduced bacterial counts at the wound site (Fig. 6a), outperforming the PL-5 group. *In vivo* antimicrobial efficacy further revealed its clinical translation potential. Besides, the inhibitory activity of R2 against fungal strains was assessed, revealing suppression of C. tropicalis growth at 32 *µg/ml* and C. albicans growth at 64 *µg/ml* with greater efficacy than PL-5. This suggests that, besides its antibacterial effects, R2 is also effective against other pathogenic microorganisms.

**Table 1.**
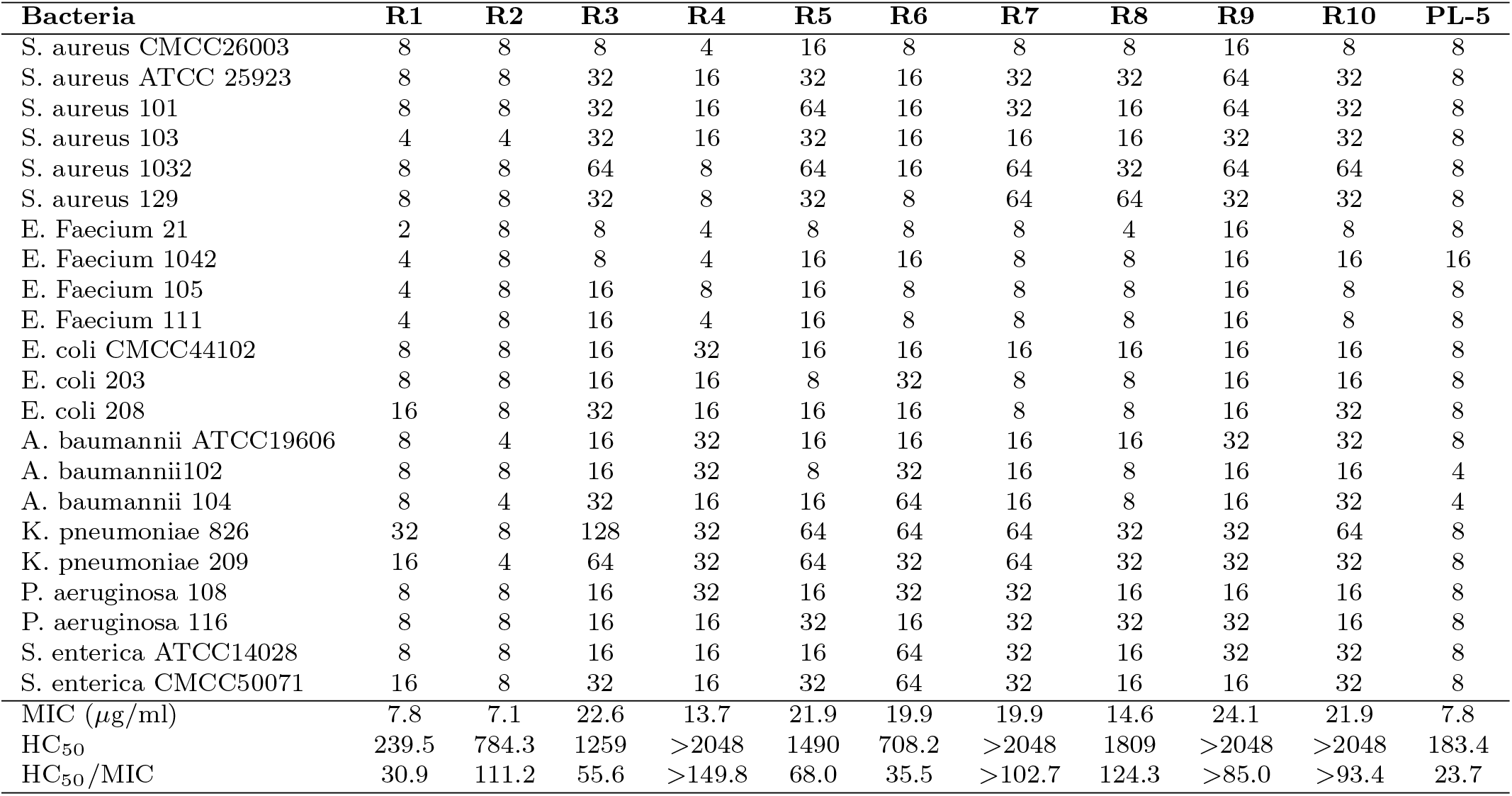
Antimicrobial activity and toxicity data of our synthetic peptides from R1 to R10. Eight out of ten of our peptides’ treatment windows are larger than PL-5.

| **Bacteria** | **R1** | **R2** | **R3** | **R4** | **R5** | **R6** | **R7** | **R8** | **R9** | **R10** | **PL-5** |
| --- | --- | --- | --- | --- | --- | --- | --- | --- | --- | --- | --- |
| *S. aureus* CMCC26003 | 8 | 8 | 8 | 4 | 16 | 8 | 8 | 8 | 16 | 8 | 8 |
| *S. aureus* ATCC 25923 | 8 | 8 | 32 | 16 | 32 | 16 | 32 | 32 | 64 | 32 | 8 |
| *S. aureus* 101 | 8 | 8 | 32 | 16 | 64 | 16 | 32 | 16 | 64 | 32 | 8 |
| *S. aureus* 103 | 4 | 4 | 32 | 16 | 32 | 16 | 16 | 16 | 32 | 32 | 8 |
| *S. aureus* 1032 | 8 | 8 | 64 | 8 | 64 | 16 | 64 | 32 | 64 | 64 | 8 |
| *S. aureus* 129 | 8 | 8 | 32 | 8 | 32 | 8 | 64 | 64 | 32 | 32 | 8 |
| *E. Faecium* 21 | 2 | 8 | 8 | 4 | 8 | 8 | 8 | 4 | 16 | 8 | 8 |
| *E. Faecium* 1042 | 4 | 8 | 8 | 4 | 16 | 16 | 8 | 8 | 16 | 16 | 16 |
| *E. Faecium* 105 | 4 | 8 | 16 | 8 | 16 | 8 | 8 | 8 | 16 | 8 | 8 |
| *E. Faecium* 111 | 4 | 8 | 16 | 4 | 16 | 8 | 8 | 8 | 16 | 8 | 8 |
| *E. coli* CMCC44102 | 8 | 8 | 16 | 32 | 16 | 16 | 16 | 16 | 16 | 16 | 8 |
| *E. coli* 203 | 8 | 8 | 16 | 16 | 8 | 32 | 8 | 8 | 16 | 16 | 8 |
| *E. coli* 208 | 16 | 8 | 32 | 16 | 16 | 16 | 8 | 8 | 16 | 32 | 8 |
| *A. baumannii* ATCC19606 | 8 | 4 | 16 | 32 | 16 | 16 | 16 | 16 | 32 | 32 | 8 |
| *A. baumannii* 102 | 8 | 8 | 16 | 32 | 8 | 32 | 16 | 8 | 16 | 16 | 4 |
| *A. baumannii* 104 | 8 | 4 | 32 | 16 | 16 | 64 | 16 | 8 | 16 | 32 | 4 |
| *K. pneumoniae* 826 | 32 | 8 | 128 | 32 | 64 | 64 | 64 | 32 | 32 | 64 | 8 |
| *K. pneumoniae* 209 | 16 | 4 | 64 | 32 | 64 | 32 | 64 | 32 | 32 | 32 | 8 |
| *P. aeruginosa* 108 | 8 | 8 | 16 | 32 | 16 | 32 | 32 | 16 | 16 | 16 | 8 |
| *P. aeruginosa* 116 | 8 | 8 | 16 | 16 | 32 | 16 | 32 | 32 | 32 | 16 | 8 |
| *S. enterica* ATCC14028 | 8 | 8 | 16 | 16 | 16 | 64 | 32 | 16 | 32 | 32 | 8 |
| *S. enterica* CMCC50071 | 16 | 8 | 32 | 16 | 32 | 64 | 32 | 16 | 16 | 32 | 8 |
| MIC ($\mu\text{g/ml}$) | 7.8 | 7.1 | 22.6 | 13.7 | 21.9 | 19.9 | 19.9 | 14.6 | 24.1 | 21.9 | 7.8 |
| HC<sub>50</sub> | 239.5 | 784.3 | 1259 | >2048 | 1490 | 708.2 | >2048 | 1809 | >2048 | >2048 | 183.4 |
| HC<sub>50</sub>/MIC | 30.9 | 111.2 | 55.6 | >149.8 | 68.0 | 35.5 | >102.7 | 124.3 | >85.0 | >93.4 | 23.7 |

**Table 2.** Antimicrobial activities, anti-fungi activities and toxicities of our discovered R2 against the other peptides.

| **Bacteria** | **R2** | **PL-5** | **C16G2** | **D2A21** | **Omiganan** | **OP-145** | **DPK-060** | **QLX-227-1** |
| --- | --- | --- | --- | --- | --- | --- | --- | --- |
| *S. aureus* CMCC26003 | 8 | 8 | 8 | 8 | 8 | 16 | 32 | 8 |
| *S. aureus* ATCC 25923 | 8 | 8 | 32 | 8 | 16 | 32 | 128 | 64 |
| *S. aureus* 101 | 8 | 8 | 32 | 8 | 16 | 32 | 128 | 64 |
| *S. aureus* 103 | 4 | 8 | 16 | 8 | 16 | 32 | 64 | 64 |
| *S. aureus* 1032 | 8 | 8 | 32 | 8 | 16 | 64 | 128 | 32 |
| *S. aureus* 129 | 8 | 8 | 32 | 8 | 16 | 32 | 128 | 64 |
| *E. Faecium* 21 | 8 | 8 | 16 | 8 | 8 | 8 | 32 | 16 |
| *E. Faecium* 1042 | 8 | 16 | 16 | 8 | 8 | 8 | 64 | 32 |
| *E. Faecium* 105 | 8 | 8 | 8 | 8 | 8 | 8 | 64 | 16 |
| *E. Faecium* 111 | 8 | 8 | 8 | 8 | 8 | 4 | 32 | 16 |
| *E. coli* CMCC44102 | 8 | 8 | 8 | 16 | 16 | 32 | — | 16 |
| *E. coli* 203 | 8 | 8 | 8 | 8 | 16 | 32 | 256 | 16 |
| *E. coli* 208 | 8 | 8 | 16 | 16 | 16 | 64 | — | 32 |
| *A. baumannii* ATCC19606 | 4 | 8 | 8 | 16 | 32 | 32 | 32 | 16 |
| *A. baumannii* 102 | 8 | 4 | 8 | 16 | 16 | 32 | 32 | 16 |
| *A. baumannii* 104 | 4 | 4 | 8 | 16 | 16 | 32 | 128 | 16 |
| *K. pneumoniae* 826 | 8 | 8 | 16 | 16 | 32 | 64 | — | 16 |
| *K. pneumoniae* 209 | 4 | 8 | 16 | 16 | 32 | 64 | — | 32 |
| *P. aeruginosa* 108 | 8 | 8 | 32 | 16 | 32 | 32 | 256 | 128 |
| *P. aeruginosa* 116 | 8 | 8 | 32 | 16 | 16 | 32 | 256 | 32 |
| *S. enterica* ATCC14028 | 8 | 8 | 8 | 8 | 32 | 32 | — | 32 |
| *S. enterica* CMCC50071 | 8 | 8 | 16 | 16 | 32 | 32 | — | 64 |
| MIC ($\mu\text{g/ml}$) | 7.1 | 7.8 | 14.6 | 11.0 | 16.5 | 26.5 | 83.0 | 28.2 |
| HC<sub>50</sub> | 784.3 | 183.4 | 2048.0 | 269.1 | 490.8 | >2048 | >2048 | 767.8 |
| IC<sub>50</sub> | 83 | 32.6 | — | — | — | — | — | — |
| HC<sub>50</sub>/MIC | 111.2 | 23.7 | >140.7 | 24.5 | 29.7 | >77.3 | >24.7 | 27.2 |
| IC<sub>50</sub>/MIC | 11.8 | 4.2 | — | — | — | — | — | — |
| **Fungi** |  |  |  |  |  |  |  |  |
| *C. tropicalis* CT1 | 32 | 64 |  |  |  |  |  |  |
| *C. albicans* 9241 | 64 | 128 |  |  |  |  |  |  |
| *C. albicans* 9113 | 64 | 128 |  |  |  |  |  |  |
| MIC ($\mu\text{g/ml}$) | 50.8 | 101.6 | | | | | | |

### Drug resistance assay

Since the mechanism of action of antimicrobial peptides does not involve specific molecular targets, the likelihood of bacteria developing resistance is relatively low [52]. To assess resistance development, we cultured *S. aureus* CMCC26003 in sub-MIC concentrations of R2, using clindamycin as a control. After 20 passages, *S. aureus* did not develop significant resistance to R2, whereas the clindamycin-treated group exhibited pronounced resistance after just five passages, which further intensified after 20 passages (Fig. 6b). The results on antimicrobial testing against multidrug-resistant strains indicate that R2 has exceptional potential for combating rapid resistance development and treating multidrug-resistant infections, laying a solid foundation for its clinical application.

### In vitro evaluation of antimicrobial safety

A major challenge in the clinical translation of cationic antimicrobial peptides is their non-specific toxicity. To evaluate the safety profile of R1-R10, we conducted cytotoxicity and hemolysis assays and compared the results with PL-5. Mouse embryonic fibroblast cells NIH/3T3 was used for cytotoxicity; R2 demonstrated higher biocompatibility than PI-5 (Table 2, and Fig. 6c and d). PL-5 exhibited higher toxicity against red blood cells, whereas R2 showed significantly reduced toxicity, indicating better blood biocompatibility (Table 2, and Fig. 6e and f). Moreover, the 50% inhibitory concentration (*IC*_50_) (Fig. 6d) and 50% hemolytic concentration (*HC*_50_) (Fig. 6f) of R2 were significantly higher than its MIC, suggesting a broader therapeutic window.

### Antimicrobial mechanism

Disruption of bacterial membranes is widely recognized as a key mechanism of action for cationic antimicrobial peptides [53]. Bacterial membranes, enriched with anionic lipid components such as lipopolysaccharides in Gram-negative bacteria or teichoic acids in Gram-positive bacteria, exhibit a significantly higher negative charge density than mammalian cell membranes, rendering them more susceptible to cationic compounds. Cationic peptides interact electrostatically with bacterial membranes, leading to pore formation, leakage of intracellular contents, and, ultimately, bacterial death. We hypothesize that R2 exerts its antimicrobial effects primarily through this mechanism. To verify this, we used scanning electron microscopy (SEM) to observe bacterial cells treated with R2. The results (Fig. 6h) revealed severe disruption of bacterial membrane integrity, further supporting the hypothesis of pore-formation-mediated action. Syto9/PI dual fluorescence staining was conducted on AMP-R2-treated *S to confirm membrane permeabilization further. aureus*. Under confocal laser fluorescence microscopy (CLSM) analysis (Fig. 6g), Syto9 (green) stains live cells with intact membranes, while PI (red) selectively enters cells with compromised membranes. In the R2-treated group, the significant red fluorescence indicated dead cells with compromised membranes. In untreated control groups, cells exhibited strong green fluorescence with minimal PI uptake. These findings confirm that R2 exhibits significantly enhanced membrane permeabilization. QLX-227-1 requires a higher concentration to effectively induce membrane permeability in bacteria cells, highlighting the superior membrane-disrupting capability of R2 compared to the source peptides.

## Discussion

We first investigate the cooperation of LLM and specialized deep learning-based models in scientific discovery by applying functional peptide recognition and generation. We show that the LLM can be engaged everywhere in conventional deep learning-based models: from the model level, the LLM can choose to ensemble which model based on their validation performance; from the data level, the LLM can provide more reasonable and precise outcome or generate the training data. By taking advantage of existing common LLM rather than training from scratch, we do not need to use thousands of the most advanced GPUs like H100 [31], which are incredibly costly to the academic circle. They also provide unique generalizations and a reasonable ability to handle unseen situations. The specialized models are designed to cater to expert domains, offering finer control and much lower training cost (*<*5,000 $) compared to LLM (*>*5,500,000 $). The specialized small models are also suitable for the scientific field with limited data since fewer parameters indicate less tendency to overfit. Besides, since we use existing LLM and train small deep learning models, all the computational tasks in this work can be finished in only one day. The updating process of the specialized small models also only takes hours. The specialized model is also more flexible than the LLM. For example, we can replace the anti-bacterial reward function with an anti-fungi reward function to generate the peptide’s target fungi. Meanwhile, additional data can quickly retrain the specialized model, thus constructing a loop of computing, wet-lab validation, and adding wet-lab data for further computing, which can significantly improve the probability of finding clinical drugs.

The out-of-domain issue has been widely discussed in the field of medical imaging analysis [54–56], but merely discussed except by [21] in the file of peptide analysis. Two typical out-of-domain cases significantly hurt the efficiency of the silicon-aided peptide discovery. One is when the sequence similarity between the training and validation sets differs. Another is that the specialized deep learning-based model can not work on unseen tasks. Our proposed PepSeek combines the extensive generalization and reasoning capabilities of LLM with the precision and flexibility of specialized small models for the first time. The specialized deep-learning-based model can reduce the hallucination of LLM. At the same time, the LLM can make the specialized deeplearning model better work on unseen or unusual data with its reasoning ability.

Our proposed PepSeek harnesses the extensive generalization and reasoning capabilities inherent to LLMs alongside the precision and flexibility afforded by specialized small models, thereby creating a unified framework that not only advances the discovery and optimization of antimicrobial peptides but also holds significant promise for broader applications in biological molecule design; by integrating disparate sources of information such as predicted protein structures [38, 49] based on the graph representation [38, 47, 57, 58], the approach further refines the accuracy of drug discovery, and with the incorporation of prompt optimization techniques like TextGrad [59], PepSeek exemplifies a trans-formative strategy that bridges the gap between heuristic generalization and targeted, precision-based analysis—a synergy that is anticipated to spur substantial progress across various domains of biomedical research.

In vitro and in vivo studies comprehensively validated the potent functionality of the novel antimicrobial peptide R2. R2 exhibited vigorous antimicrobial activity against a broad spectrum of clinically relevant pathogens by employing membrane permeabilization as its primary mechanism. Compared to antimicrobial peptides currently in clinical trials, R2 demonstrated equal or superior efficacy while offering enhanced safety through its excellent cellular compatibility, resulting in a significantly expanded therapeutic index. However, future in vivo safety and stability evaluations are also necessary to broaden the application of R2 [60]. Notably, R2 posed a minimal risk of inducing resistance, addressing a critical challenge in combating antibiotic resistance. In a full-thickness skin excision infection mouse model, R2 achieved therapeutic outcomes significantly superior to PL-5, highlighting its translational potential for infection treatment. In addition, although our model is not fungus-specific, antimicrobial experiments validated the antifungal ability of R2, and we can turn our pipeline to fungus-targeted by replacing the reward function in PRL. The case study in Table 2 also validates that PepSeek can provide successful zero-shot predictions. These findings underscore the promise of integrating LLMs and deep learning models to accelerate the discovery of antibiotic peptide. Furthermore, PepSeek shows a versatile and user-friendly platform for de novo design and optimization of therapeutic peptides, facilitates rapid adaptation to diverse research needs.

## Methods

### Dataset

We compiled our dataset by integrating peptides from multiple sources to capture a breadth of activity while mitigating potential issues inherent to data scarcity. Functional peptide data were drawn from DBASSP [39] and QLAPD [38], and for negative samples, we utilized entries from UniProt (http://www.uniprot.org). In contrast to previous studies that classified peptides as bioactive or non-bioactive, we adopted the methodology described in [38] by categorizing peptides according to their minimum inhibitory concentration (MIC). Specifically, peptides with a MIC below 128 were deemed active, while negative peptides were assigned a MIC value of 512. This approach allowed us to more precisely distinguish peptides based on their functional potency and toxicity profiles. There are 26,565 AMPs for training and validation in total.

To ensure robustness and minimize data leakage, we cleaned the merged dataset comprehensively and implemented a five-fold cross-validation strategy. Peptides exhibiting more than 30% internal similarity (as determined by the Needleman– Wunsch algorithm [40]) were removed from overlapping folds to simulate out-of-domain scenarios. In addition to cross-validation, the model was tested on an independent set of private 151 synthetic peptide sequences [38]. To further address the challenge of limited data on peptides with less common functions (e.g., antiviral activity), we focused primarily on antibacterial peptides during training and evaluation while also leveraging the zero-shot predictive capabilities of the LLM for novel function detection on datasets such as dbAMP 3.0 [37] that cover a wider range of peptide functions.

### Implementation and Evaluation

All models were trained for 120 epochs, and test performance was determined by averaging the performance metrics from the final epoch across five folds. For the classification task, we evaluated model performance using the area under the receiver operating characteristic curve (AUROC) and the area under the precision-recall curve (AUPRC). For regression, we employed mean absolute error (MAE), relative squared error (RSE), Pearson correlation coefficient (PCC), and Kendall correlation coefficient (KCC). Our computational infrastructure comprises an Intel Core i9-14900K CPU, two NVIDIA 4090D GPU with 24GB of VRAM, and 192GB of RAM. Storage is provided by a Samsung 990 PRO SSD (4TB), and all experiments were executed on Ubuntu 22.04 with NVIDIA Driver Version 550.90.07 and CUDA Version 12.4. The software stack includes PyTorch [61] v2.2.0, Python 3.10.0, Biopython [62] v1.78 for obtaining global sequence attributes, and torchmetrics [42] v1.4.0 for metric evaluation.

The following equations define the evaluation metrics. For classification, given true positives (*TP*), false positives (*FP*), false negatives (*FN*), and true negatives (*TN*), we define the 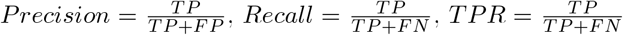, and 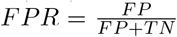. The area under the precision-recall curve (AUPRC) is given by:

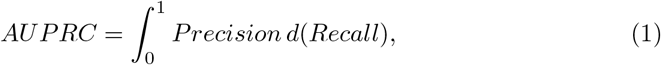

and the area under the receiver operating characteristic curve (AUROC) is defined as

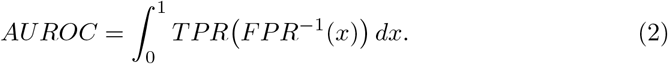

For regression, the mean absolute error (MAE) is defined as

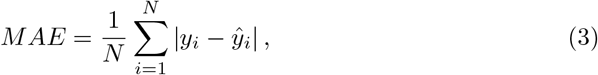

the relative squared error (RSE) as

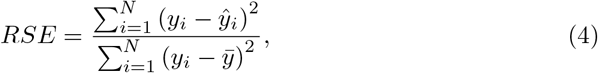

the Pearson correlation coefficient (PCC) as

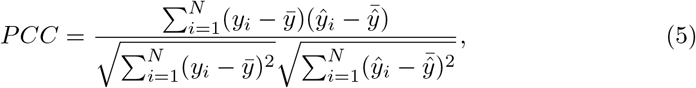

and the Kendall correlation coefficient (KCC) [41] as

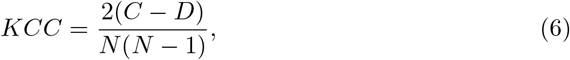

where *C* and *D* denote the numbers of concordant and discordant pairs, respectively.

### Construction of Specialized Deep Learning Models for Peptide Identification and Regression

To mitigate potential overfitting associated with models possessing many parameters, such as ProGen [19], we adopted several advanced architectures from previous studies [22]. We evaluated them alongside conventional models for both peptide classification and regression tasks. In our implementation, we focused on four representative models:

1. **LSTM:** We implemented a long short-term memory network [25] that processes peptide sequences by capturing both short-term and long-term dependencies. Its architecture includes standard gates (input, forget, and output) that regulate the cell state and hidden state updates.
2. **Transformer and Multi-head Attention (MHA):** Our Transformer-based [26] model leverages self-attention mechanisms to encode peptide sequences. Multi-head Attention enables the model to capture diverse aspects of the sequence by simultaneously attending to different subspaces of the input features.
3. **Multi-head Latent Attention (MLA):** This model extends the standard MHA [33] by incorporating a latent attention component that integrates global sequence information. The latent attention assists in capturing hidden dependencies and enhances performance on complex sequence data.
4. **Mamba:** The Mamba model [44] relies on a convolutional feature extractor and an attention-based refinement module. This architecture effectively combines local feature extraction with global contextualization, facilitating more robust peptide property prediction.

We employ a weighted cross-entropy loss function to handle class imbalance for the peptide classification task. Given *N* training samples and *C* classes, the loss is defined as:

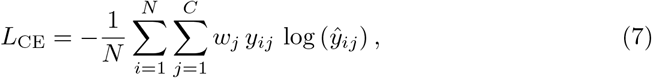

where *y*_*ij*_ is the binary ground-truth indicator for class *j* of sample 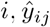 is the predicted probability for class *j*, and *w*_*j*_ is the corresponding class weight.

For the regression task, we utilize the mean squared error (MSE) loss, defined as:

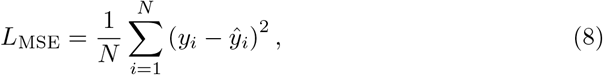

where *y*_*i*_ is the true value (e.g., MIC) for sample *i* and prediction 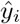 is the corresponding

In contrast to earlier approaches that employed a straightforward ensemble of various models, we leveraged a LLM (deepseek-r1) to intelligently select and ensemble a minimal subset of models based on their validation performance. This LLM-guided ensemble strategy dynamically weights the outputs of the chosen models, achieving comparable or superior performance while reducing the overall model complexity. The ensemble output is computed as follows:

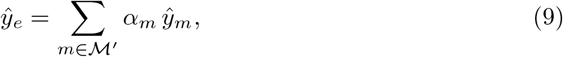

where ℳ^*′*^denotes the subset of selected models and *α*_*m*_ represents the weight assigned to model *m*.

### Design of Prior-Guided Reinforcement Learning for Peptide Generation and Optimization

#### State Space Design

In this study, the state space, 𝒮, was meticulously constructed to incorporate both the primary sequence (**Seq**) and its associated physicochemical properties **Phy**. Specifically, the state comprises the sequence (e.g., “KWMIKWP-SNWFTML” from [38]), key properties such as the Helix Fraction, and net charge at pH 7, in addition to the current mutation position (**p**). This comprehensive representation ensures that the reinforcement learning (RL) algorithm can make decisions informed by both sequence and property data [48]. The state space is defined as

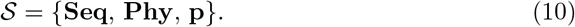

#### Action Space Design

The action space, 𝒜, is a 20-dimensional discrete set representing the standard amino acids. This design accommodates both the necessity for mutation and the restrictions based on the properties of the current amino acid (*AA*):

1. Mandatory Mutation for Specific Amino Acids: If *AA* belongs to a predefined set of undesirable amino acids, bad aa = {*A, E, D, C, N, Q, M, H*}, a mutation must be induced towards any amino acid not included in bad aa.
2. Restrictions Based on Properties: If *AA* is positively charged or *AA* ∈ {*I, V, L*}, all mutation actions are masked except for the no mutation action (*a*_0_), effectively preserving the amino acid. All other residues are allowed if mutations do not result in an amino acid from bad aa.

The discrete action space is expressed as

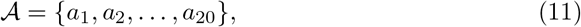

The mutation mask is defined by

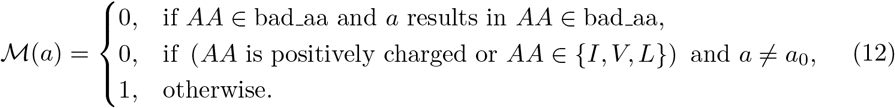

#### Reward Function Design

To quantitatively assess the quality of the generated sequences, we define a composite reward function that evaluates six physicochemical properties: the *helix fraction* (*H*), the *isoelectric point* (*pI*), the *net charge at pH 7* (*C*), the *positive charge ratio* (*P*), the *hydrophobic ratio* (*R*), and the *instability index* (*I*). For each property, an ideal target range and an optimal value are specified as follows:

- **Helix Fraction:** Target range [3, 6] with optimal value 4.5.
- **Isoelectric Point:** Target range [0.6, 0.9] with optimal value 0.75.
- **Net Charge at pH 7:** Target range [1, 5] with optimal value 3.
- **Positive Charge Ratio:** Target range [0.1, 0.3] with optimal value 0.2.
- **Hydrophobic Ratio:** Target range [0.3, 0.6] with optimal value 0.45.
- **Instability Index:** The desired value is below 1 (the lower, the better).

##### Intermediate Reward Calculation

For each property, if the current value *x* lies within its target range and has an optimal value 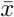 with tolerance Δ, the intermediate reward is calculated by

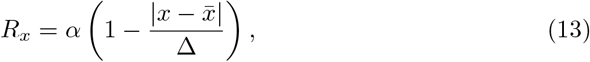

where the coefficient is set to *α* = 0.3. The tolerances for properties are given by:

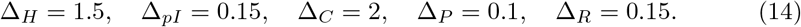

If a property falls outside its specified range, a − *α* penalty is applied. For example, the reward for the helix fraction is:

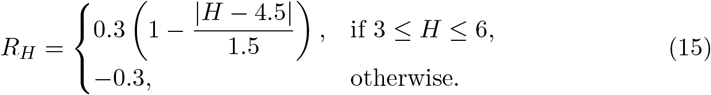

A similar formulation applies to *pI, C, P*, and *R*. For the instability index, which is desired to be below 1, the reward is given by

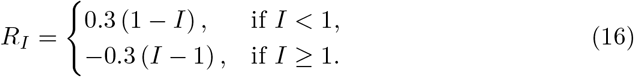

*Final Reward and Overall Evaluation:* Once sequence generation is complete, a final evaluation is performed. If all properties satisfy their criteria:

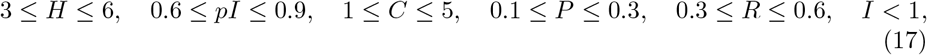

an additional reward of +5 is added; otherwise, a penalty of − 5 is imposed. Let the cumulative reward from all intermediate calculations be *R*. With the final adjustment, we have

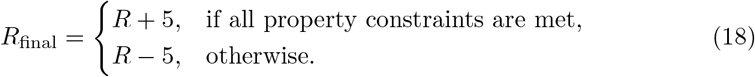

Furthermore, if the mutation number *m* exceeds a predetermined threshold *m*_max_, an extra penalty of −5 is incurred:

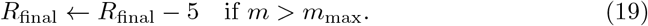

In addition to the above reward components, specialized deep learning models are employed to predict further properties of the peptide, such as antibacterial activity, toxicity, and the minimum inhibitory concentration (MIC). The outputs of these models contribute additional reward terms.

For peptide classification, let *p*_anti_ be the predicted probability (from a deep learning classifier) that the peptide exhibits antibacterial activity, and let *p*_toxic_ be the probability that the peptide is hemolytic toxic. A probability-based scaling factor *λ*_proba_ is constructed based on the selectivity ratio (*HC*_50_ */ MIC*).

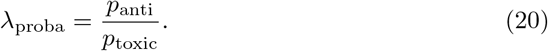

A deep learning model predicts peptide regression’s minimum inhibitory concentration (MIC). Assuming that a lower MIC indicates higher efficacy, the regression reward term is defined as

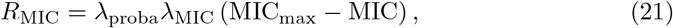

where MIC_max_ denotes the reference maximum value in the prediction space, and *λ*_MIC_ is a weighting factor.

The overall total reward is then computed as the sum of the previously defined property-based reward *R*_final_ and the deep learning model rewards:

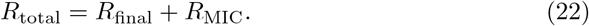

This comprehensive reward strategy allows the evaluation framework to assess sequence quality by integrating structural and chemical properties with the predictions of specialized deep-learning models, thus aligning with the objectives of generating peptides with desired biological activities. Implementing the Proximal Policy Optimization [63] algorithm follows established guidelines to ensure effective policy updates. This includes employing clipping to maintain the ratio of new to old policies within a reasonable range, using multiple epochs of stochastic gradient descent per data sample, and employing separate networks or network heads for the policy and value functions to stabilize learning. This approach helps manage the exploration-exploitation balance more effectively and adapt the policy to complex environments such as those encountered in peptide sequence optimization [64].

#### Candidate Sequence Generation and Filtering with PRL

State inputs were constructed for all valid positions within a given sequence to generate candidate sequences. The action distribution is computed for each position, and the mutation probability, *P*, is derived. This probability, defined as

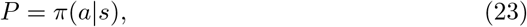

reflects the exploration efficiency of the RL model, with higher values indicating sequences likely to yield elevated rewards [65, 66]. Given that the RL-generated peptides may be various, a ranked list of candidate sequences is generated. Subsequent filtering further refines these candidates by prioritizing peptides with high antimicrobial activity and low toxicity.

### Setting of Using Large Language Models

In this study, we developed a comprehensive approach for peptide design by integrating LLMs with specialized small models. Considering that the best open-sourced LLM is DeepSeek-R1 [31], we use it in our cooperation. DeepSeek-R1 also provides a much lower token fee and transparent reasoning process, making it better for downstream analysis. For comparison, we use the deepseekv3 [33], gpt4o [30], and o3-mini [34]. The detailed prompts are shown in Fig. 5.

### Materials and Bacterial strains

Peptides with a purity of *>* 95% were custom-synthesized by GenScript. Apexbio supplied the Cell Counting Kit-8 (CCK-8). Invitrogen supplied the LIVE/DEAD BacLight bacterial viability kit. DMEM medium and fetal bovine serum were purchased from ZhongQiaoXinZhou and Abbkine. The bacterial strains used in this study are listed in Supplementary Fig. 7, which includes detailed strain information. Standard bacterial strains were purchased from the China Medical Culture Collection (CMCC) and China Center of Industrial Culture Collection (CICC). Multi-drug-resistant bacterial strains and Candida albicans were clinically isolated at Qilu Hospital of Shandong University.

### Minimum inhibitory concentration

The peptides’ minimum inhibitory concentration (MIC) against bacterial and fungal strains was determined using the microdilution method following the Clinical and Laboratory Standards Institute (CLSI) guidelines. For bacterial strains, peptide solutions (0.5–256 *µg/ml*) were prepared via twofold serial dilutions in Mueller-Hinton broth (MHB) and mixed with 50 *µL* of bacterial suspension (1 × 10^6^ CFU/ml) in a 96-well plate. The plate was incubated at 37°C for 18 hours. MIC determination followed the CLSI M27-A3 guidelines for fungal strains using the micro broth dilution method. Strains were cultured on YPD agar plates and incubated at 30°C overnight, then transferred to YPD liquid medium and shaken at 200 rpm until the mid-logarithmic phase was reached. The fungal suspension (1 × 10^3^ cells/mL) was prepared in Mops-buffered RPMI 1640 medium (pH 7.4), and 100 *µL* was added to each well of a 96-well plate containing a peptide concentration gradient (0.5–256 *µg/ml*). Fluconazole served as a positive control, and untreated wells were negative controls. After incubation at 35°C for 24 hours, the MIC was defined as the lowest peptide concentration that completely inhibited fungal growth.

### Mechanism of action: membrane disruption

The membrane permeability changes induced by AMPs were analyzed using SEM and TEM. *S. aureus* CMCC26003 was cultured at 37°C overnight to the exponential phase. Bacterial suspensions (10^8^ CFU/ml) were treated with R2 at a final concentration of 1 × MIC for 3 h at 37°C, with untreated cells as the control. Samples were washed three times with PBS and fixed with glutaraldehyde overnight. The sample was sequentially dehydrated with ethanol/water solutions of different concentrations. The specimens were then observed using a SEM (HITACHI S-4700). To further assess its effects on bacterial membrane integrity, Syto9/PI fluorescence staining was conducted on *S. aureus* cells treated with 1× MIC of AMP-R2 for 3 hours.

### Longitudinal resistance development compared to Clindamycin

To assess the potential for resistance development, *S. aureus* CMCC26003 was subjected to serial passaging in sub-MIC concentrations of AMP-R2 and Clindamycin over 20 days. During the experiment, bacterial cultures were exposed daily to fresh media containing the respective compounds at sub-MIC levels, with MIC values measured at regular intervals to monitor changes in susceptibility.

### Analysis of cytotoxicity

To evaluate the cytotoxicity of AMP-R2, the NIH/3T3 cell line was used. For the cell line, 8 × 10^3^ cells were seeded into individual wells of 96-well plates and cultured at 37°C with 5% CO_2_ overnight to allow cell attachment. The following day, the culture medium was carefully removed, and fresh media containing AMP-R2 at the desired concentrations was added to each well. The cells were incubated with AMP-R2 for 24 hours under standard culture conditions. After the treatment, the media were replaced with fresh DMEM containing 10% CCK-8 reagent, followed by an additional 1–2 hours incubation at 37°C. The absorbance of the solution in each well was measured at 450 nm using a microplate reader. The percentage of cell viability was calculated using the following formula:

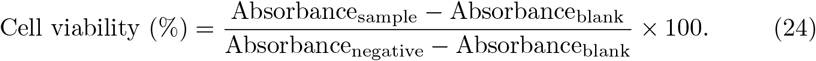

### Evaluation of hemolytic activity

The hemolytic activity of the peptides was evaluated using red blood cells (RBCs). Fresh mice blood containing an anticoagulant (0.25% sodium citrate) was centrifuged at 1000 × *g* for 5 minutes to collect the RBCs. The RBCs were washed three times with sterile PBS and then diluted to an 8% (v/v) concentration. The absorbance at 570 nm was measured using a microplate reader. RBCs treated with PBS were used as the negative control (OD_0_), while RBCs treated with 1% Triton X-100 (v/v) were used as the positive control (OD_100_). The minimum hemolytic concentration (MHC) was defined as the peptide concentration that caused 10% hemolysis of RBCs. The percentage of hemolysis was calculated using the following formula:

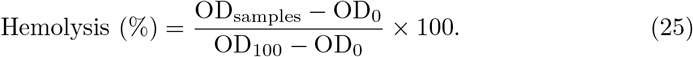

### In vivo therapeutic efficacy in a full-thickness skin wound infection model

Female C57BL/6J mice (7 − 8 weeks) were used in a mouse skin infection model. The mice were purchased from Beijing Vital River Laboratory Animal Technology Co., Ltd. All animal experiments were reviewed and approved by the Ethics Committee for Laboratory Animals of Qilu Hospital of Shandong University (DWLL-2023-104). To assess the therapeutic efficacy of AMP-R2 in vivo, a full-thickness skin wound infection model was established in mice using *S. aureus* CMCC26003. Specifically, after hair removal from the back of the mice, full-thickness wounds with a diameter of 5 mm were created on the dorsal skin and infected with 5 *µ*l of bacterial suspension at a concentration of 10^8^ CFU/ml. Four hours post-infection, the treatment group received a topical application of 10 *µ*l of AMP-R2 peptide solution (10 mg/ml). PL-5 (Peceleganan) was used as a positive control (10 mg/ml). Skin samples from the infected wounds were collected 24 hours later for bacterial quantification.

## Data availability

The data related to this work is available in at https://drive.google.com/drive/folders/1nDLcBYnFyey71BUcHqdKRG3O1jJrKFY?usp=sharing.

## Code availability

The source codes and trained models are accessible in the supplementary material. All this information will be made publicly available after acceptance.

## Acknowledgments

We thank the Translational Medicine Core Facility of Shandong University for consulting and lending instruments that supported this work. The scientific calculations in this study were performed on the HPC Cloud Platform of Shandong University. This work was financially supported by the National Natural Science Foundation of China (Grant No. 82270580 and 82470571) and the Taishan Scholars Program of Shandong Province to YL; National Natural Science Foundation Young Student Basic Research Project of China (Grant No. 824B2012) to YW.

## Author Contributions

Conceptualization: YL, HG, YW, QK, XL; Methodology: HG, YW, QK, XL; Investigation: HG, YW, QK, XL, LL, BW, YZ, JZ, GC, JC, YY, XY, XZ; Visualization: HG, QK, XL, BW; Funding acquisition: YL; Project administration: YL, LL; Supervision: 19 YL, XZ, XY, YY; Writing – original draft: HG, QK, XL, BW; Writing – review & editing: YL, YW, HG, QK.

## Supplementary Materials

**Fig. 1.**
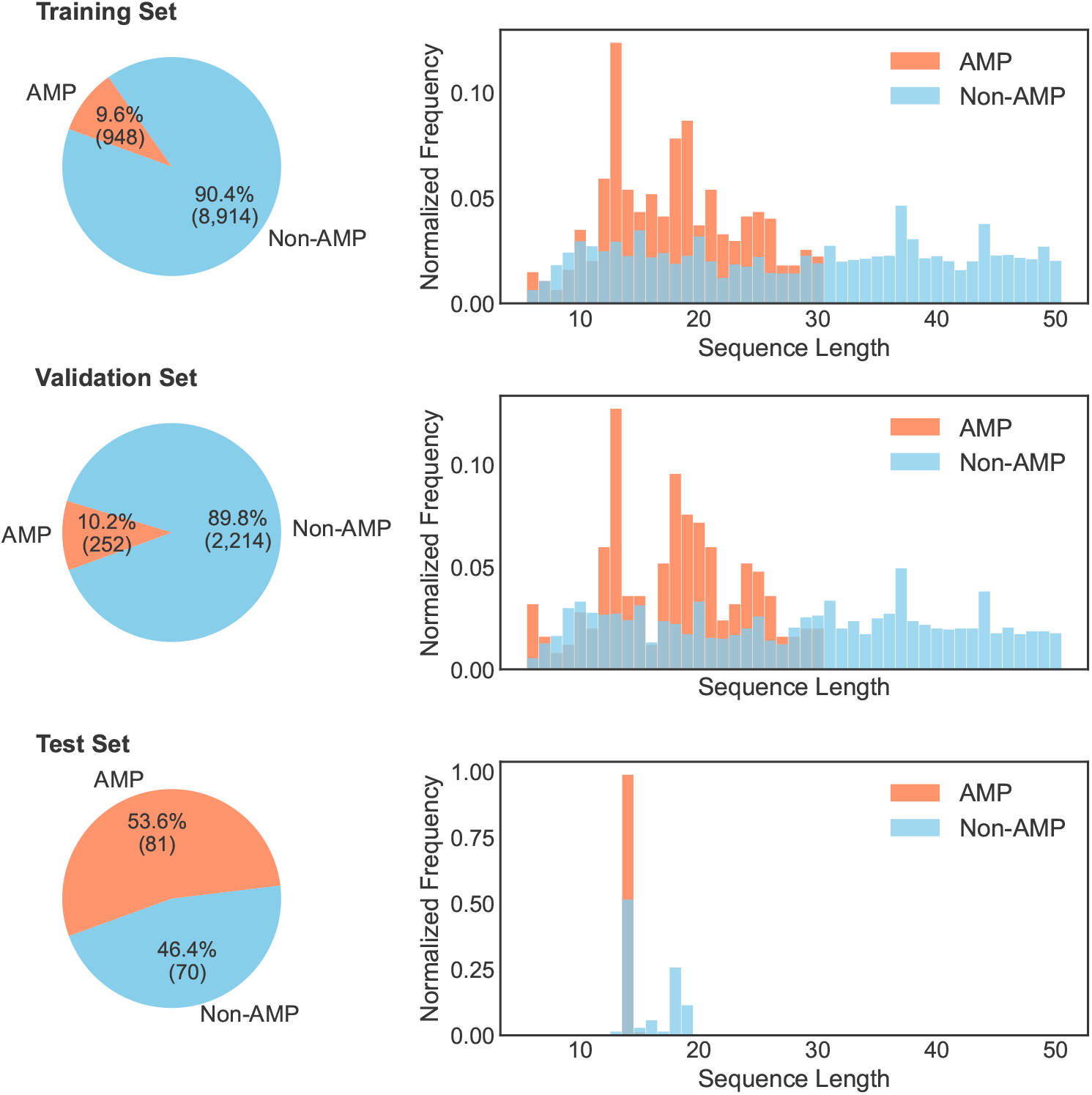
Data distribution for the AMP classification task. To emulate a challenging real-world scenario, the testing dataset exhibits a distribution shift relative to the source domain. Moreover, employing a lower threshold for peptide identification leads to a pronounced imbalance between the training and validation sets, further compounding the challenge.

**Fig. 2.**
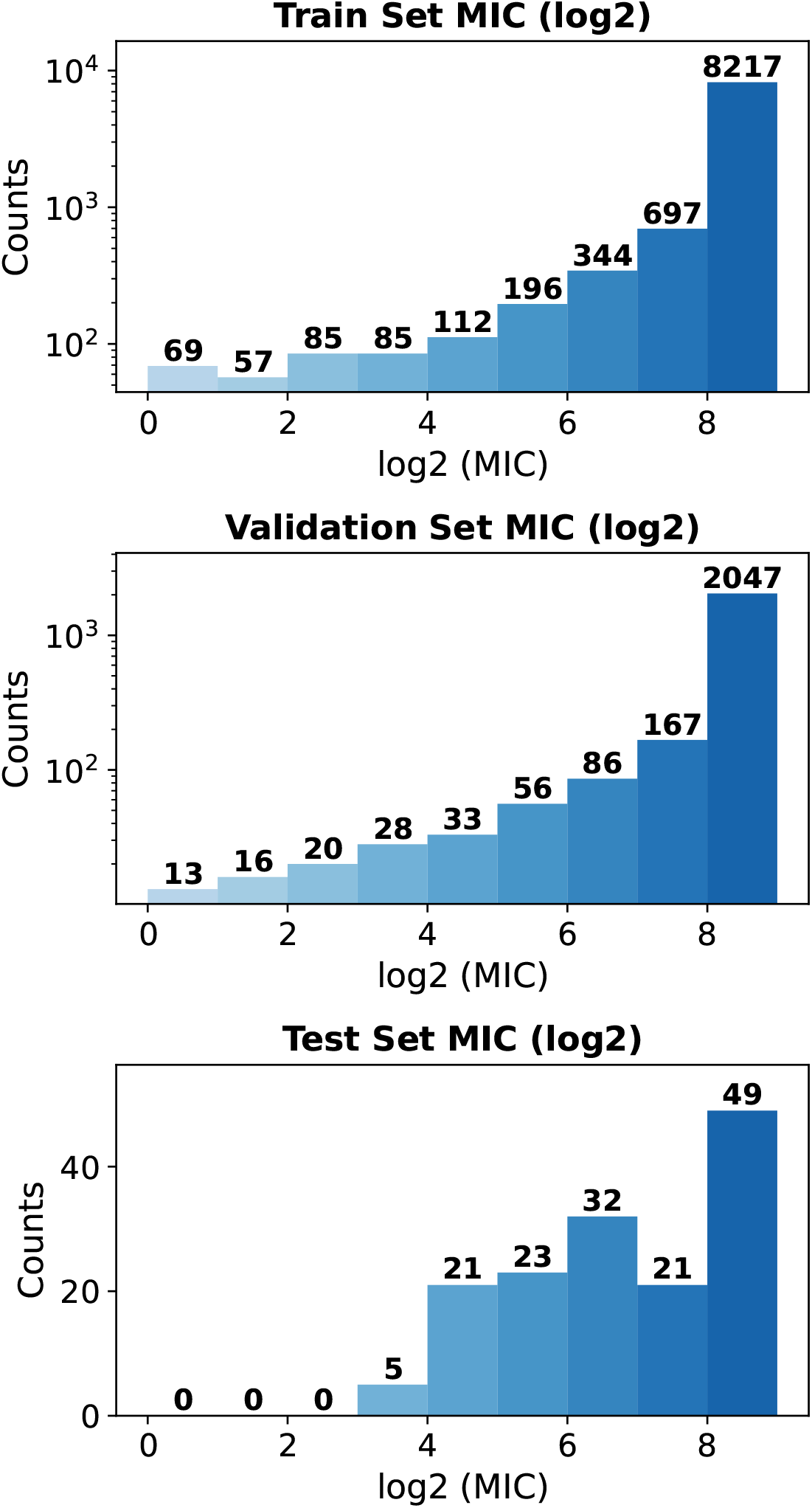
Data distribution for the AMP MIC regression task. The MIC values follow a longtailed distribution with fewer samples at the lower end, posing a challenge for robust identification. Notably, the test set was derived from rigorous wet-lab assessments—averaging results from over ten bacterial species—resulting in a scarcity of peptides with extremely low MIC values.

**Fig. 3.**
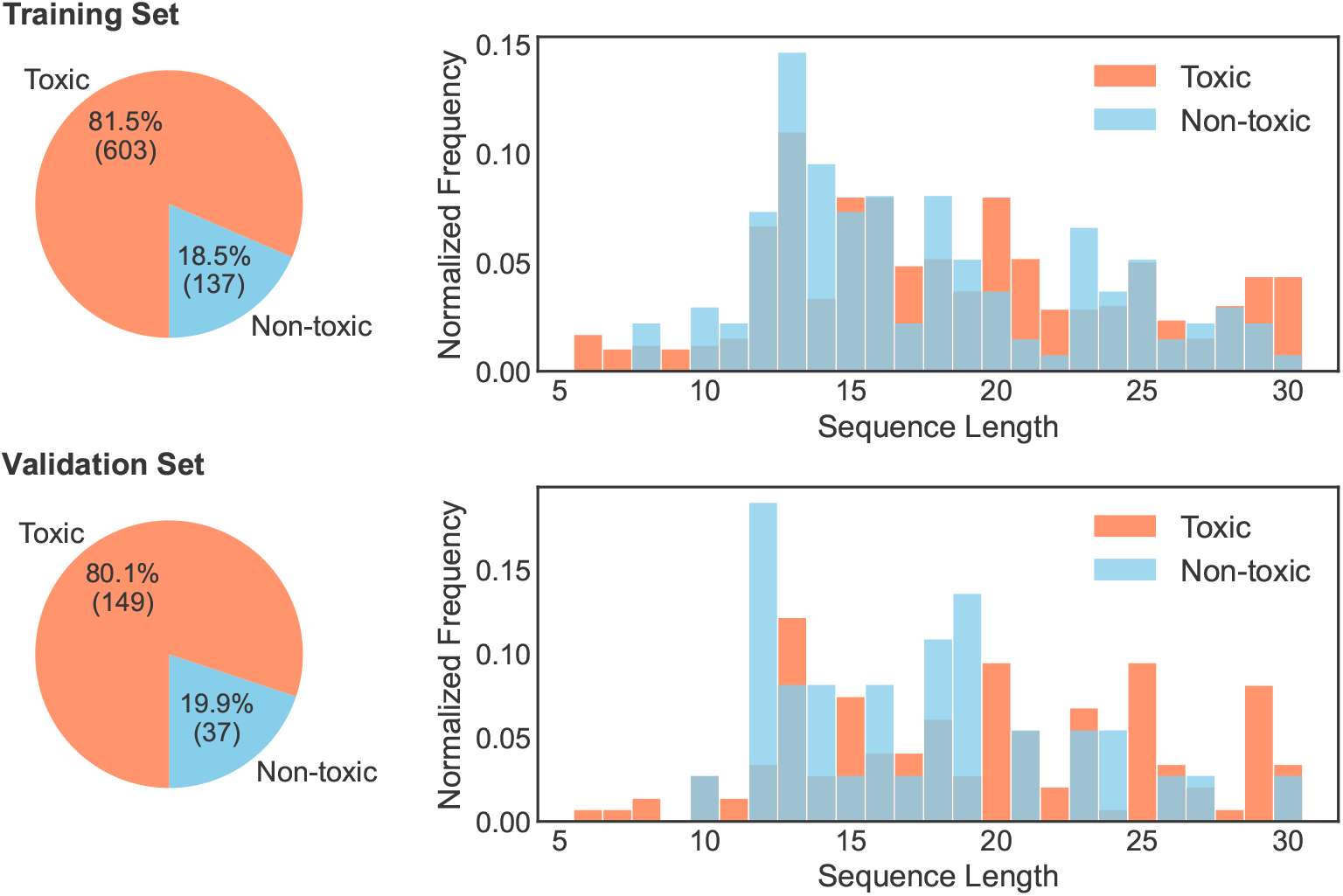
Data distribution for the hemolytic toxic classification task. To ensure safety, a high threshold for peptide identification also leads to a pronounced imbalance between the training and validation sets, compounding the challenge.

**Fig. 4.**
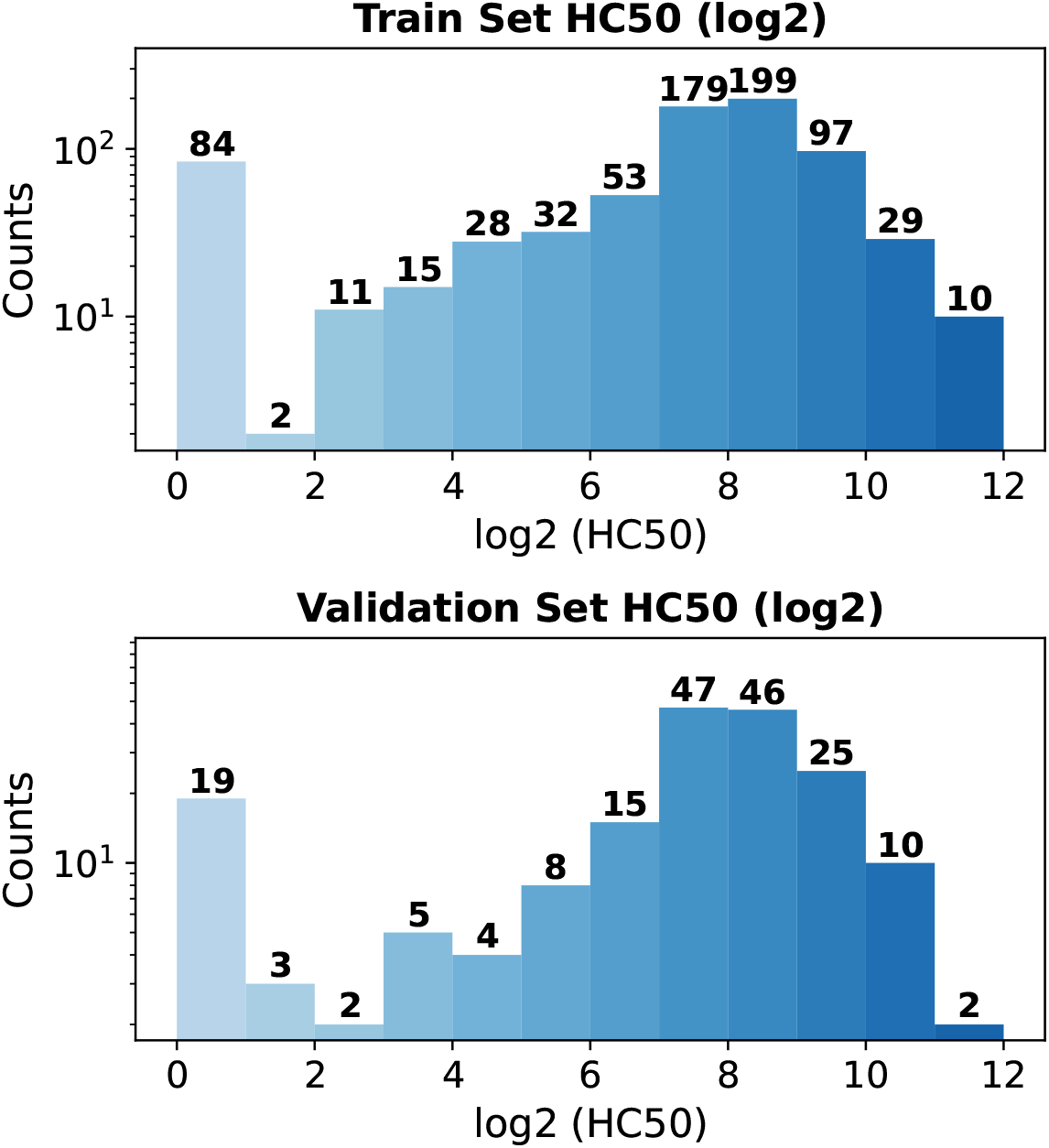
Data distribution for the HC50 regression task. The HC50 values follow a near Gaussian distribution with fewer samples toward both ends and a spike at the low end, posing a challenge for robust identification.

**Fig. 5.**
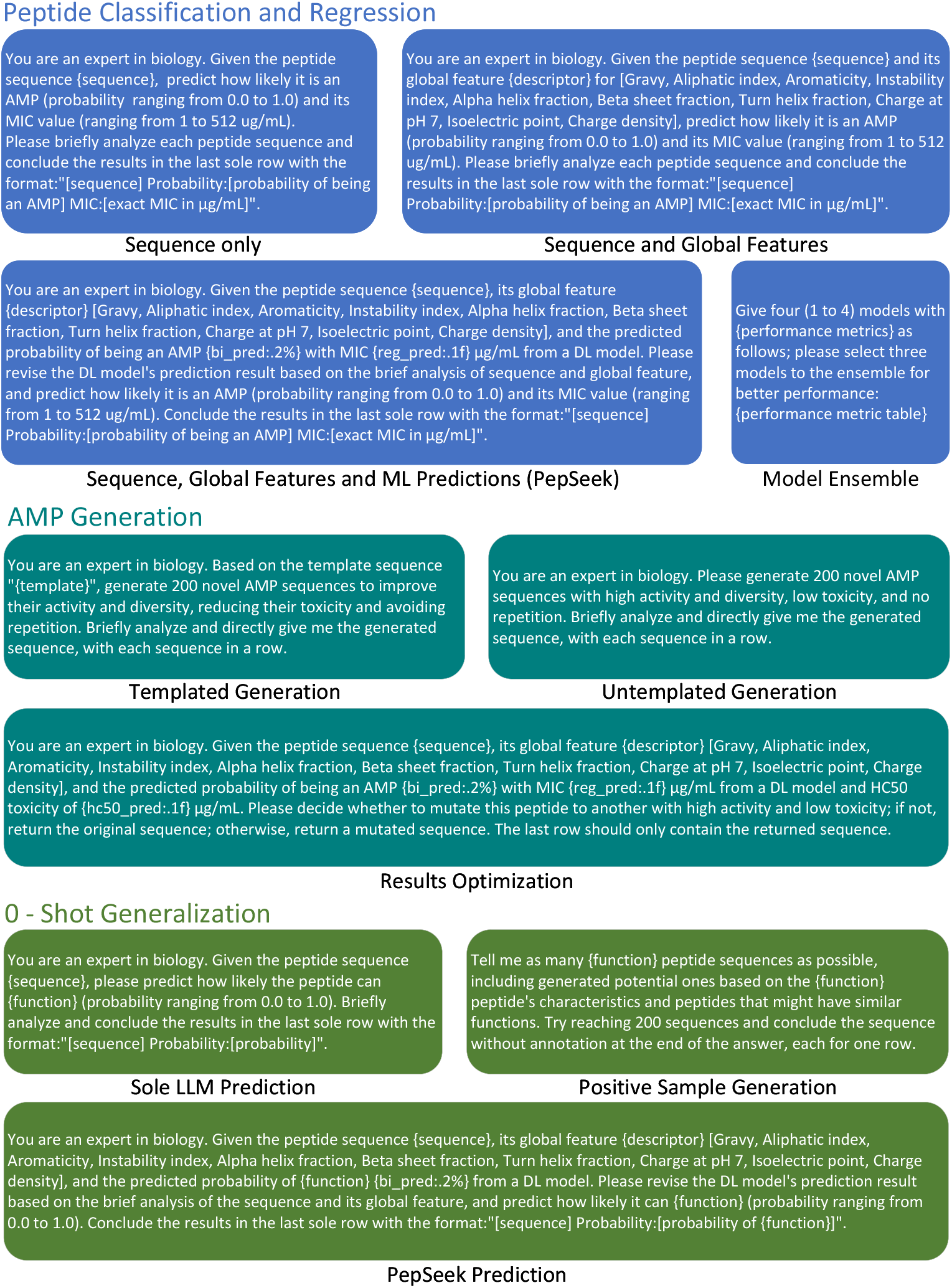
Large-language model (LLM) prompt templates employed in this study. Prompts are categorized into recognition, generation, and zero-shot generalization.

**Fig. 6.**
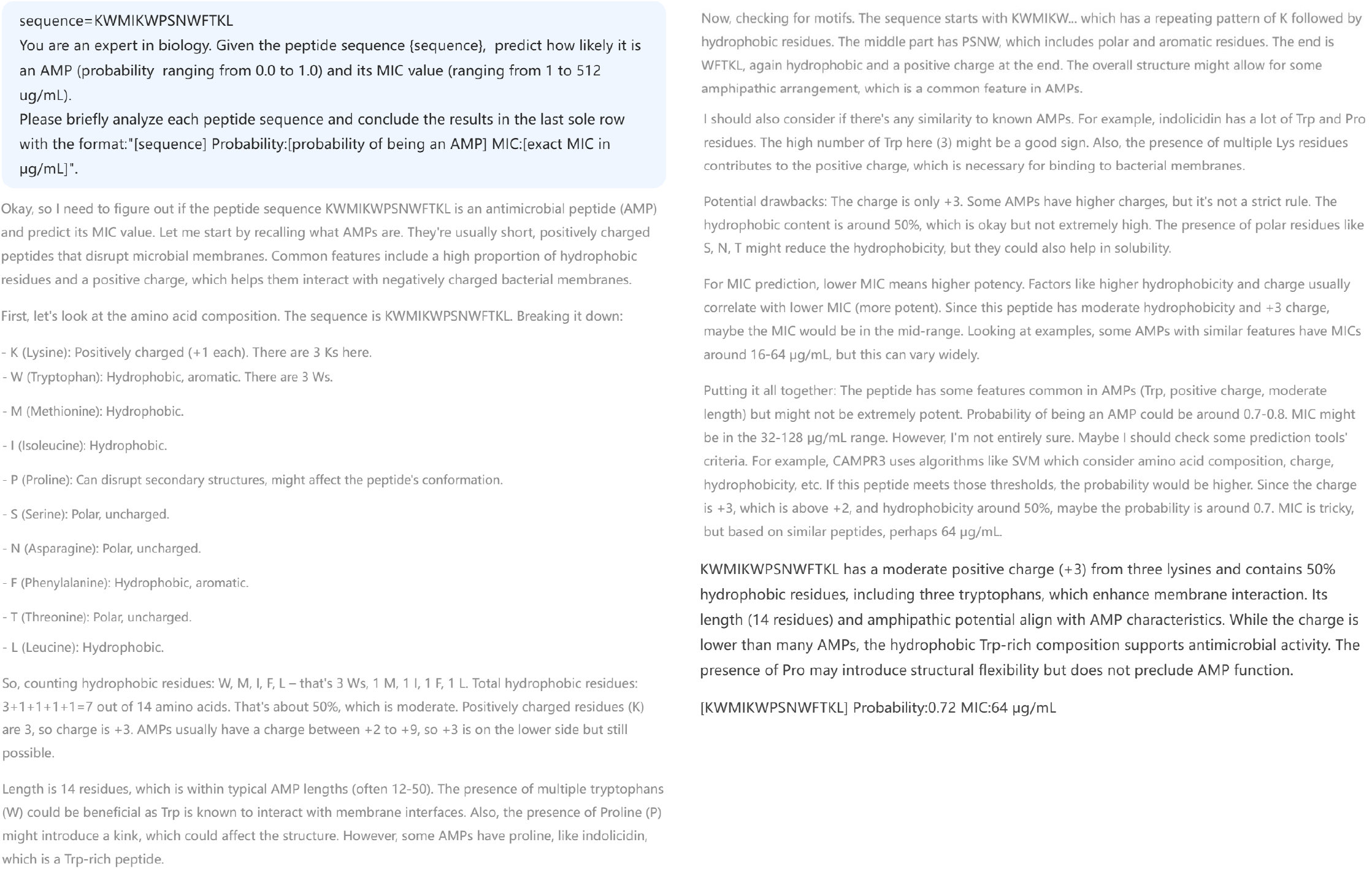
Visualization of the reasoning process of the LLM (DeepSeek-R1 [31]). The LLM demonstrates the capacity to infer peptide function from amino acid composition, accurately compute overall net charge, and iteratively refine its analysis to deliver a detailed, critical assessment.

**Fig. 7.**
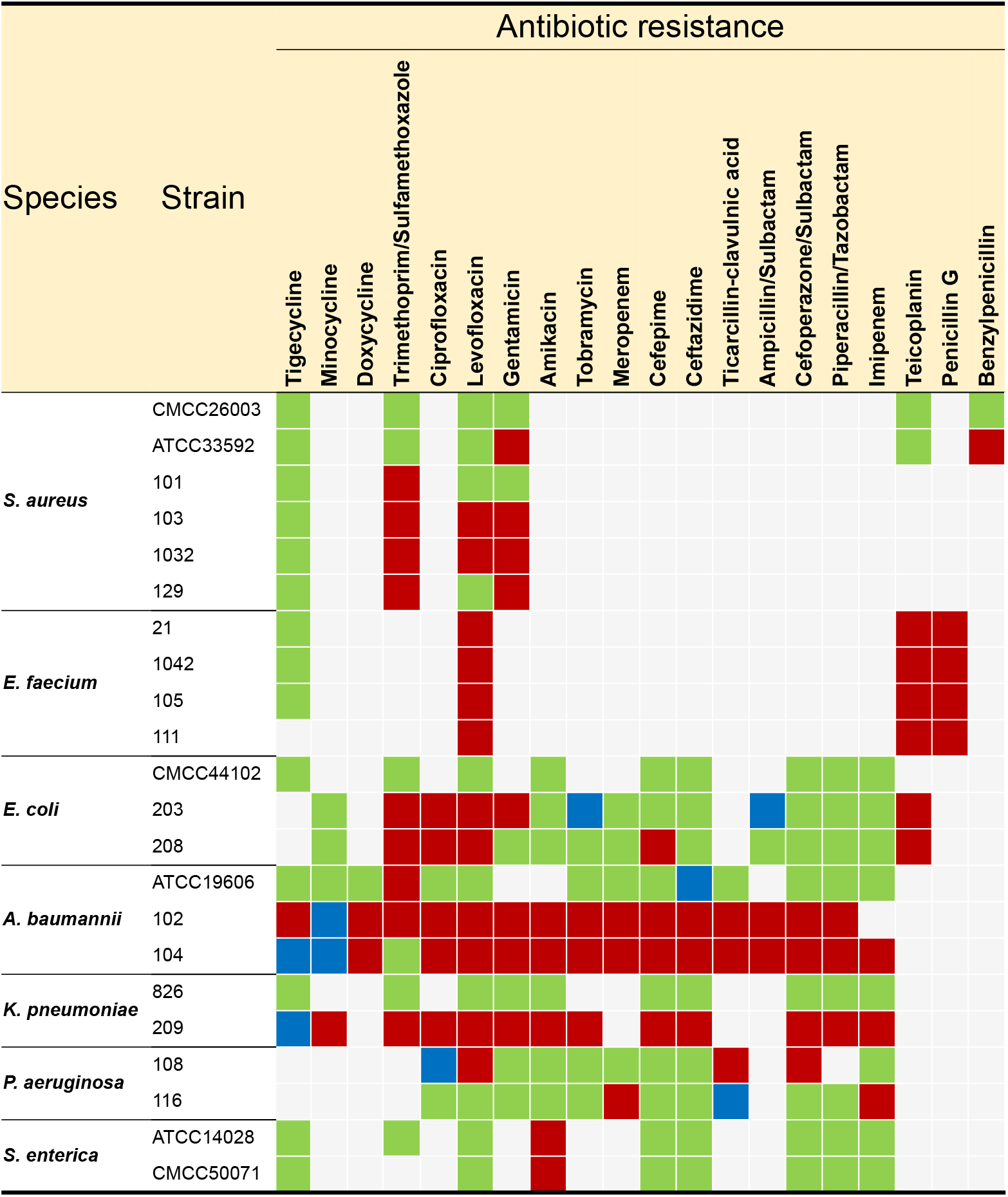
Bacterial strain information. Green, red, and blue boxes denote clinically isolated strains that are susceptible, resistant, and moderately susceptible to the corresponding antibiotics, respectively; gray boxes indicate that susceptibility to the respective agents was not evaluated.

**Table 1.**
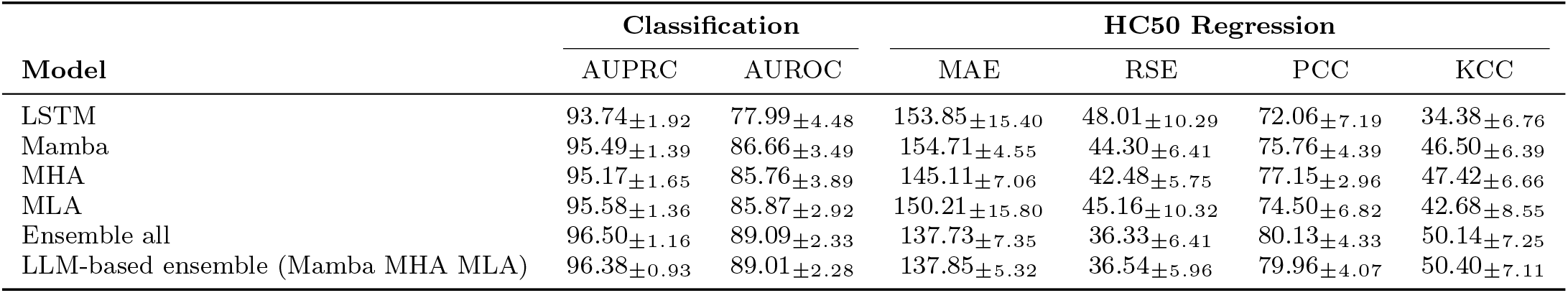
Model Performance Comparison on Hemolytic Toxin Classification and HC50 Regression Tasks. Comparison of model performance on hemolytic toxin classification and HC50 regression tasks. The cooperative ensemble strategy effectively identifies configurations that perform as well as or better than the individual small models.

**Table 2.**
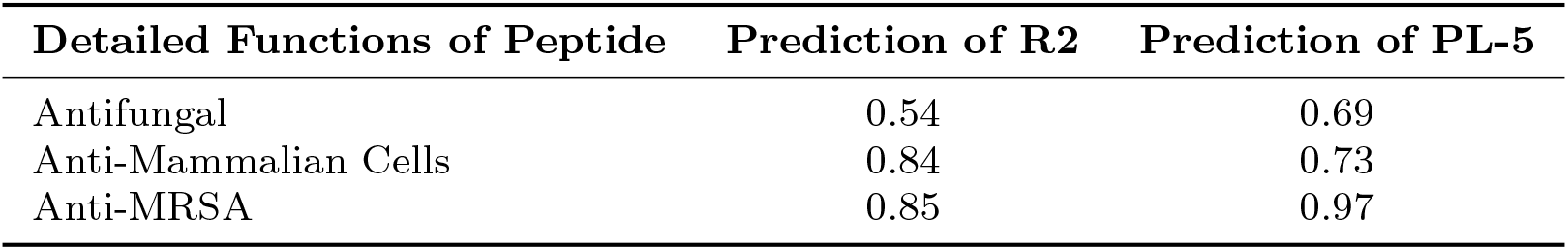
Zero-shot prediction of peptide functionalities. Predictions from PeekSeek for zero-shot recognition (refer to Section 2.3) indicate that peptides R2 and PL-5 exhibit significant antifungal, anti-mammalian cell, and anti-MRSA activities. These computational insights are consistent with the corresponding wet lab results reported in Table 2.

## References

[1] Zhao, T. et al. A foundation model for joint segmentation, detection and recognition of biomedical objects across nine modalities. Nature Methods 1–11 (2024).

[2] Li, T. et al. A foundation model identifies broad-spectrum antimicrobial peptides against drug-resistant bacterial infection. Nature Communications 15, 7538 (2024).

[3] Moor, M. et al. Foundation models for generalist medical artificial intelligence. Nature 616, 259–265 (2023).

[4] Cui, H. et al. scgpt: toward building a foundation model for single-cell multi-omics using generative ai. Nature Methods 1–11 (2024).

[5] Wang, X. et al. A pathology foundation model for cancer diagnosis and prognosis prediction. Nature 634, 970–978 (2024).

[6] Peng, C., Shang, J., Guan, J., Wang, D. & Sun, Y. Viralm: empowering virus discovery through the genome foundation model. Bioinformatics 40, btae704 (2024).

[7] Ren, X. et al. Hydrogelfinder: A foundation model for efficient self-assembling peptide discovery guided by non-peptidal small molecules. Advanced Science 2400829 (2024).

[8] Dong, C. et al. A survey of natural language generation. ACM Computing Surveys 55, 1–38 (2022).

[9] Achiam, J. et al. Gpt-4 technical report. arXiv preprint arXiv:2303.08774 (2023).

[10] Huang, Z. et al. A pathologist–ai collaboration framework for enhancing diagnostic accuracies and efficiencies. Nature Biomedical Engineering 1–16 (2024).

[11] Hou, W. & Ji, Z. Assessing gpt-4 for cell type annotation in single-cell rna-seq analysis. Nature Methods 1–4 (2024).

[12] Rehana, H. et al. Evaluating gpt and bert models for protein–protein interaction identification in biomedical text. Bioinformatics Advances 4, vbae133 (2024).

[13] Orsi, M. & Reymond, J.-L. Can large language models predict antimicrobial peptide activity and toxicity? RSC Medicinal Chemistry (2024).

[14] Chen, Y. & Zou, J. Simple and effective embedding model for single-cell biology built from chatgpt. Nature Biomedical Engineering 1–11 (2024).

[15] Huang, Z., Bianchi, F., Yuksekgonul, M., Montine, T. J. & Zou, J. A visual– language foundation model for pathology image analysis using medical twitter. Nature medicine 29, 2307–2316 (2023).

[16] Simon, E., Swanson, K. & Zou, J. Language models for biological research: a primer. Nature Methods 21, 1422–1429 (2024).

[17] Wan, F., Wong, F., Collins, J. J. & de la Fuente-Nunez, C. Machine learning for antimicrobial peptide identification and design. Nature Reviews Bioengineering 2, 392–407 (2024).

[18] Kaplan, J. et al. Scaling laws for neural language models. arXiv preprint arXiv:2001.08361 (2020).

[19] Madani, A. et al. Large language models generate functional protein sequences across diverse families. Nature Biotechnology 41, 1099–1106 (2023).

[20] Ma, Y. et al. Identification of antimicrobial peptides from the human gut microbiome using deep learning. Nature Biotechnology 40, 921–931 (2022).

[21] Huang, J. et al. Identification of potent antimicrobial peptides via a machine-learning pipeline that mines the entire space of peptide sequences. Nature Biomedical Engineering 7, 797–810 (2023).

[22] Wang, B. et al. Explainable deep learning and virtual evolution identifies antimicrobial peptides with activity against multidrug-resistant human pathogens. Nature Microbiology 1–16 (2025).

[23] Chen, T. & Guestrin, C. Xgboost: A scalable tree boosting system, 785–794 (ACM, 2016).

[24] Prokhorenkova, L., Gusev, D., Vorobev, A., Dorogush, A. & Gulin, E. Catboost: unbiased boosting with categorical features, 6638–6648 (2018).

[25] Hochreiter, S. & Schmidhuber, J. Long short-term memory. Neural Computation 9, 1735–1780 (1997).

[26] Vaswani, A. et al. Guyon, I. et al. (eds) Attention is all you need. (eds Guyon, I. et al.) Advances in Neural Information Processing Systems 30: Annual Conference on Neural Information Processing Systems 2017, December 4-9, 2017, Long Beach, CA, USA, 5998–6008 (2017).

[27] Ben-David, S., Blitzer, J., Crammer, K. & Pereira, F. Analysis of representations for domain adaptation. Advances in neural information processing systems 19 (2006).

[28] Xu, Z., Gong, H., Wan, X. & Li, H. Greenspan, H. et al. (eds) Asc: Appearance and structure consistency for unsupervised domain adaptation in fetal brain mri segmentation. (eds Greenspan, H. et al.) MEDICAL IMAGE COMPUTING AND COMPUTER ASSISTED INTERVENTION, MICCAI 2023, PT VII, Vol. 14226, 325–335 (2023).

[29] Farahani, A., Voghoei, S., Rasheed, K. & Arabnia, H. R. A brief review of domain adaptation. Advances in data science and information engineering: proceedings from ICDATA 2020 and IKE 2020 877–894 (2021).

[30] Hurst, A. et al. Gpt-4o system card. arXiv preprint arXiv:2410.21276 (2024).

[31] Guo, D. et al. Deepseek-r1: Incentivizing reasoning capability in llms via reinforcement learning. arXiv preprint arXiv:2501.12948 (2025).

[32] Wei, J. et al. Chain-of-thought prompting elicits reasoning in large language models. Advances in neural information processing systems 35, 24824–24837 (2022).

[33] Liu, A. et al. Deepseek-v3 technical report. arXiv preprint arXiv:2412.19437 (2024).

[34] Jaech, A. et al. Openai o1 system card. arXiv preprint arXiv:2412.16720 (2024).

[35] Ji, Z. et al. Survey of hallucination in natural language generation. ACM Computing Surveys 55, 1–38 (2023).

[36] Farquhar, S., Kossen, J., Kuhn, L. & Gal, Y. Detecting hallucinations in large language models using semantic entropy. Nature 630, 625–630 (2024).

[37] Yao, L. et al. dbamp 3.0: updated resource of antimicrobial activity and structural annotation of peptides in the post-pandemic era. Nucleic Acids Research 53, D364–D376 (2025).

[38] Wang, Y. et al. De novo multi-mechanism antimicrobial peptide design via multimodal deep learning. bioRxiv 2024–01 (2024).

[39] Pirtskhalava, M. et al. Dbaasp v3: database of antimicrobial/cytotoxic activity and structure of peptides as a resource for development of new therapeutics. Nucleic acids research 49, D288–D297 (2021).

[40] Needleman, S. B. & Wunsch, C. D. A general method applicable to the search for similarities in the amino acid sequence of two proteins. Journal of Molecular Biology 48, 443–453 (1970).

[41] Abdi, H. The kendall rank correlation coefficient. Encyclopedia of measurement and statistics 2, 508–510 (2007).

[42] Detlefsen, N. S. et al. Torchmetrics-measuring reproducibility in pytorch. Journal of Open Source Software 7, 4101 (2022).

[43] Nijkamp, E., Ruffolo, J. A., Weinstein, E. N., Naik, N. & Madani, A. Progen2: exploring the boundaries of protein language models. Cell systems 14, 968–978 (2023).

[44] Dao, T. & Gu, A. Transformers are ssms: Generalized models and efficient algorithms through structured state space duality ([Organization Name], 2024).

[45] Liu, A. et al. Deepseek-v2: A strong, economical, and efficient mixture-of-experts language model. arXiv preprint arXiv:2405.04434 (2024).

[46] Gong, H., Huang, R., Chen, G., Li, G. et al. Sysu-hcp at vqa-med 2021: A data-centric model with efficient training methodology for medical visual question answering., 1218–1228 (2021).

[47] Gong, H. et al. Unbiased curriculum learning enhanced global-local graph neural network for protein thermodynamic stability prediction. Bioinformatics 39, btad589 (2023).

[48] Sutton, R. S. & Barto, A. G. Reinforcement Learning: An Introduction (MIT Press, 2018).

[49] Jumper, J. et al. Highly accurate protein structure prediction with alphafold. nature 596, 583–589 (2021).

[50] Li, G., Lai, Z. & Shan, A. Advances of antimicrobial peptide-based biomaterials for the treatment of bacterial infections. Advanced Science 10, 2206602 (2023).

[51] Wei, Y. et al. Peceleganan spray for the treatment of skin wound infections: A randomized clinical trial. JAMA Network Open 7, e2415310–e2415310 (2024).

[52] Mookherjee, N., Anderson, M. A., Haagsman, H. P. & Davidson, D. J. Antimicrobial host defence peptides: functions and clinical potential. Nature reviews Drug discovery 19, 311–332 (2020).

[53] Gan, B. H., Gaynord, J., Rowe, S. M., Deingruber, T. & Spring, D. R. The multi-faceted nature of antimicrobial peptides: Current synthetic chemistry approaches and future directions. Chemical Society Reviews 50, 7820–7880 (2021).

[54] Xu, L. et al. Real-time monitoring of manual acupuncture stimulation parameters based on domain adaptive 3d hand pose estimation. Biomedical Signal Processing and Control 83, 104681 (2023).

[55] Gong, H. et al. Diffuse-uda: Addressing unsupervised domain adaptation in medical image segmentation with appearance and structure aligned diffusion models. arXiv preprint arXiv:2408.05985 (2024).

[56] Gong, H., Yu, L., Wan, X. & Li, H. Domain generalized medical landmark detection via robust boundary-aware pre-training, 1–9 (2025).

[57] Liang, B., Gong, H., Lu, L. & Xu, J. Risk stratification and pathway analysis based on graph neural network and interpretable algorithm. BMC bioinformatics 23, 394 (2022).

[58] Huang, W. et al. Bcnet: Bronchus classification via structure guided representation learning. IEEE Transactions on Medical Imaging (2024).

[59] Yuksekgonul, M. et al. Textgrad: Automatic” differentiation” via text. arXiv preprint arXiv:2406.07496 (2024).

[60] Zhu, Y. et al. Association of idealized amphiphiles and protease inhibitors: Conferring antimicrobial peptides with stable antibacterial activity under physiological conditions to combat multidrug-resistant bacteria. Drug Resistance Updates 79, 101183 (2025).

[61] Paszke, A. et al. Pytorch: An imperative style, high-performance deep learning library. Advances in neural information processing systems 32 (2019).

[62] Cock, P. J. et al. Biopython: freely available python tools for computational molecular biology and bioinformatics. Bioinformatics 25, 1422 (2009).

[63] Schulman, J., Wolski, F., Dhariwal, P., Radford, A. & Klimov, O. Proximal policy optimization algorithms. arXiv preprint arXiv:1707.06347 (2017).

[64] Huang, S. et al. The 37 implementation details of proximal policy optimization (2022). The ICLR Blog Track 2023.

[65] Kingma, D. P. & Ba, J. Adam: A method for stochastic optimization. arXiv preprint arXiv:1412.6980 (2014).

[66] Silver, D., Schrittwieser, J., Simonyan, K. & et al. Mastering the game of go without human knowledge. Nature 550, 354–359 (2020).

